# Single Nucleotide Substitutions Effectively Block Cas9 and Allow for Scarless Genome Editing in *Caenorhabditis elegans*

**DOI:** 10.1101/2021.09.14.460298

**Authors:** Jeffrey C. Medley, Shilpa Hebbar, Joel T. Sydzyik, Anna Y. Zinovyeva

**Affiliations:** Division of Biology, Kansas State University, Manhattan, Kansas, United States of America

**Author notes:** To whom correspondence should be addressed: **Corresponding Author**: Anna Zinovyeva, Mailing Address: 28 Ackert Hall, Manhattan, KS 66506.

**Keywords:** Genome Editing, CRISPR, Cas9, Blocking, Scarless, miRNA, non-coding RNA, *C. elegans*, *let-7*

## Abstract

In *Caenorhabditis elegans*, germline injection of Cas9 complexes is reliably used to achieve genome editing through homology-directed repair of Cas9-generated DNA breaks. To prevent Cas9 from targeting repaired DNA, additional blocking mutations are often incorporated into homologous repair templates. Cas9 can be blocked either by mutating the PAM sequence that is essential for Cas9 activity or by mutating the guide sequence that targets Cas9 to a specific genomic location. However, it is unclear how many nucleotides within the guide sequence should be mutated, since Cas9 can recognize “off-target” sequences that are imperfectly paired to its guide. In this study, we examined whether single-nucleotide substitutions within the guide sequence are sufficient to block Cas9 and allow for efficient genome editing. We show that a single mismatch within the guide sequence effectively blocks Cas9 and allows for recovery of edited animals. Surprisingly, we found that a low rate of edited animals can be recovered without introducing any blocking mutations, suggesting a temporal block to Cas9 activity in *C. elegans*. Furthermore, we show that the maternal genome of hermaphrodite animals is preferentially edited over the paternal genome. We demonstrate that maternally provided haplotypes can be selected using balancer chromosomes and propose a method of mutant isolation that greatly reduces screening efforts post-injection. Collectively, our findings expand the repertoire of genome editing strategies in *C. elegans* and demonstrate that extraneous blocking mutations are not required to recover edited animals when the desired mutation is located within the guide sequence.

## Introduction

The CRISPR/Cas9 system has become increasingly used to facilitate genome editing in numerous organisms (Ma and Liu, 2015; Shrock and Güell, 2017; Ma et al., 2018.). Cas9 (CRISPR associated protein 9) is a programmable endonuclease whose specificity is governed by a guide RNA that has sequence complementarity to a specific genomic location (Jinek et al., 2012). The guide RNA comprises two molecules: the CRISPR RNA (crRNA) that contains a 20-nucleotide guide sequence and a trans-acting CRISPR RNA (tracrRNA) that forms a duplex with the crRNA and bridges the guide RNA to Cas9 (Deltcheva et al., 2011; Jinek et al., 2012). A protospacer-adjacent motif (PAM) sequence is located immediately downstream of the RNA guide-complementary genomic sequence and is required for Cas9 to initiate a double-stranded DNA break. In the case of *Streptococcus pyogenes* Cas9, commonly used for genome editing, the nucleotide PAM sequence is NGG, where N is any nucleotide. (Mojica et al., 2009; Marraffini et al., 2010; Sashital et al., 2012; Jinek et al., 2012). Once a double-stranded DNA break is created, the break is typically repaired through one of two mechanisms: non-homologous end joining (NHEJ) or homology-directed repair (HDR) (Yang et al., 2020; Scully et al., 2019; Han and Huang, 2020; Li and Xu, 2016; Ceccaldi et al., 2016). In NHEJ, the broken DNA is repaired through direct ligation of the broken DNA ends. However, this process is error prone as the ligation often requires processing of the broken ends, resulting in additions or deletions of nucleotide bases at the break site (Zhao et al., 2020; Chang et al., 2017). Conversely, HDR uses a donor DNA molecule that has homology surrounding the break site as a template to precisely repair the broken DNA (Sun et al., 2020; Ranjha et al., 2018; Haber, 2018). Therefore, HDR has been widely adapted to repair Cas9-generated DNA breaks to introduce precise genome edits in a broad range of organisms. Donor repair templates can be exogenously provided as single-stranded oligodeoxynucleotides (ssODN) or double stranded DNA (dsDNA) molecules for the purpose of genome editing (Zhao et al., 2014; Yoshimi et al., 2016; Cong et al., 2013; Paquet et al., 2016; Gallagher et al., 2020). During CRISPR/Cas9-mediated genome editing, the process of HDR using a ssODN repair template is referred to as single-stranded template repair (SSTR), and results in higher genome editing efficiencies than HDR pathways that use dsDNA repair templates (Dokshin et al., 2018; Katic et al., 2015; Okamoto et al., 2019; Liu et al., 2019; Gallagher and Haber, 2021; Gallagher et al., 2020; Richardson et al., 2018).

Once a Cas9-generated DNA break is repaired through SSTR, Cas9 must be prevented from continuing to target the repaired DNA. To accomplish this, additional blocking mutations are often incorporated into homologous repair templates, disrupting the ability of Cas9 to target the repaired sequence. As the PAM is absolutely required for Cas9 activity (Mojica et al., 2009; Marraffini et al., 2010; Sashital et al., 2012; Jinek et al., 2012), the most straightforward way to block Cas9 is to introduce silent mutations into the PAM. Alternatively, Cas9 can be blocked by introducing mutations into the guide sequence, which targets Cas9 to a specific genomic location (Deltcheva et al., 2011; Jinek et al., 2012). However, studies in human cells have shown that Cas9 is capable of recognizing off-target sequences that are imperfectly paired to its guide RNA (Jinek et al., 2012; Pattanayak et al., 2013; Jiang and Doudna, 2015). Mismatches near the 3’ end of the guide RNA appear to be more effective at blocking Cas9 compared to mismatches toward the 5’ end of the guide RNA (Jinek et al., 2012; Fu et al., 2013; Pattanayak et al., 2013; Hsu et al., 2013; Mali et al., 2013; Cong et al., 2013; Zhang et al., 2015). Increasing the number of mismatches generally leads to increased blocking efficacy (Jinek et al., 2012; Fu et al., 2013; Pattanayak et al., 2013; Hsu et al., 2013; Mali et al., 2013; Cong et al., 2013; Zhang et al., 2015). Nevertheless, Cas9 has been reported to cleave DNA sequences containing up to five mismatches to certain guide RNAs (Hsu et al., 2013), although three mismatches effectively block Cas9 for most guide RNAs (Fu et al., 2013; Hsu et al., 2013; Mali et al., 2013). Therefore, it remains unclear how many nucleotides should be mismatched and where the mismatches should be located within the guide sequence to effectively block Cas9 for genome editing *in vivo*.

In the nematode *Caenorhabditis elegans*, injection of Cas9 ribonucleotide protein (RNP) complexes and ssODN repair templates into the germline of hermaphrodite animals has been reliably used to facilitate heritable genome editing (Paix et al., 2015; Paix et al., 2017; Farboud et al., 2018). However, certain types of genome edits remain challenging to design due to the need to block Cas9 from targeting the repaired DNA. Protein coding sequences are highly amenable for genome editing experiments, as codon redundancy frequently allows silent blocking mutations to be introduced without changing the amino acid identity. However, genome editing of regulatory and non-protein coding portions of the genome remain a challenge. It is often difficult to predict how extraneous blocking mutations may affect the function of non-coding regulatory sequences such as non-coding RNAs, untranslated regions, and other regulatory elements. Extraneous blocking mutations can be avoided when the intended edit also alters a PAM site and eliminates Cas9 ability to recut a repaired genome site. In such cases, genome editing is performed in a ‘scarless’ fashion. However, the dinucleotide GG of the PAM sequence (NGG) is only expected to occur, on average, every 16 bases and must overlap with the desired edit to generate a scarless edit. This frequency is likely reduced in non-coding regions that are often AT-rich. It has been suggested that single nucleotide substitutions located within three nucleotides of the PAM are sufficient to allow for genome editing in *C. elegans* (Paix et al., 2017; Farboud et al., 2018). This is of particular interest for scarless genome editing, as non-PAM mutations could block Cas9 and thereby bypass the need for additional blocking mutations. We reasoned that single nucleotide substitutions beyond the three PAM-adjacent nucleotides, located in the guide-binding region could effectively block Cas9, further facilitating scarless genome editing of non-coding sequences. Toward this end, we have performed a systematic analysis of the blocking efficacy of single nucleotide mismatches throughout the guide sequence in *C. elegans*. We demonstrate that single nucleotide substitutions throughout the guide-binding sequence are sufficient to block Cas9 and allow for effective recovery of edited animals. Strikingly, we were able to recover heritable genome edited strains without introducing any blocking mutations, suggesting that a temporal block to Cas9 activity limits the ability of Cas9 to target repaired DNA. We also show that editing of the maternal genome of self-fertile hermaphrodite animals occurs at much greater frequency compared to editing of the paternal genome. Finally, we propose a new method of mutant isolation that selects for maternally provided haplotypes and greatly reduces screening efforts post-injection. As a proof-of-principle, we use this method to generate otherwise scarless genome edits in the *let-7* microRNA. Our collective findings expand the repertoire of possible genome edits in *C. elegans* and will facilitate scarless editing of non-coding sequences.

## Materials and Methods

### *C. elegans* Strains and Genetics

All *C. elegans* strains were derived from the wild-type N2 strain and maintained at 20°C unless otherwise noted. Strains were grown under standard conditions using Nematode Growth Medium (NGM) plates seeded with *Escherichia coli* OP50 (Brenner, 1974). A full list of strains used in this study is provided in Table S1.

For experiments examining maternal vs. paternal editing, matings were performed by adding several wild-type males to a plate containing L4-staged, *tra-2* mutant hermaphrodite animals generated in this study. For *let-7* genome editing experiments, several males containing the *tmc24* balancer were mated to L4-staged, wild-type hermaphrodite animals. Animals were allowed to mate 16-24h prior to injection.

Successful mating was verified by genotyping F1 animals through *Rsa*I digestion to confirm *tra-2* heterozygosity, or presence of pharyngeal Venus to confirm presence of the *tmc24* balancer.

### CRISPR/Cas9 Genome Editing

Commercially available *S. pyogenes* Cas9 (IDT, Alt-R® S.p. Cas9 Nuclease V3) was injected at a final concentration of 2.65 µM and was included in all injection mixes. All injection mixtures also contained 200 µM KCI and 7.5 µM HEPES [pH 7.4]. Each injection mix contained one or two crRNAs targeting *dpy-10* (5 µM), *tra-2* (50µM) or *let-7* (50µM), and equimolar tracrRNA was included (5µM, 50µM or 55 µM). Single-stranded DNA oligonucleotides for *dpy-10* (3µM), *tra-2* (6µM) and *let-7 (*6µM) were used to facilitate homology-directed repair (Table S2).

To generate non-Dumpy genome edits at the *tra-2* locus, which we used to examine how effectively single nucleotide substitutions blocked Cas9, we replaced *dpy-10* co-conversion with the Pmyo-2::mCherry co-injection marker (5 ng/µL). The Pmyo2::mCherry co-injection marker was included to mark broods that were successfully injected and to enrich for genome-edits as previously described (Prior et al., 2017). A full list of oligonucleotides (IDT) used in this study is provided in Table S2.

### Screening and Genotyping

For experiments testing the efficacy of *tra-2* editing, we blindly sequenced (i.e. without *Rsa*l digestion) the F2 generation Dumpy and non-Dumpy animals originating from each F1 Roller animal. F1 Rollers were picked from jackpot broods (Paix et al., 2015) that were defined as having >20 F1 Rollers from a single P0 injected animal. F1 rollers that did not produce Dumpy, Roller and non-Dumpy progeny were excluded from our analysis. The *tra-2* genomic locus was PCR amplified using the primers 5’-CTGCTAAAGGTTAGTTGTT-3’ and 5’-ATAATGTATTCTTCATTGTTCG-3’ and sequenced using the primer 5’-ATTTTAGGAATAATTGGAGCC-3’. All *Rsa*l digestions were performed at 37°C for at least 1 h. A *Rsa*l-positive control was included on all gels used for quantification to confirm successful *Rsa*l digestion occurred.

To examine genome editing of *let-7*, we singled F1 generation Roller animals from jackpot broods that showed pharyngeal Venus signal that indicated successful mating to the tmc24 balancer pre-injection. We then blindly sequenced F2 generation animals lacking pharyngeal Venus that were therefore homozygous for the maternally provided X-chromosome haplotype. The *let-7* genomic locus was PCR amplified using the primers 5’-GTTTGCGTATGTGTATGTAG-3’ and 5’-TCCCCTGAAAATAAAACATGA-3’ and sequenced using the primer 5’-TATTCTAGATGAGTAGCCCA-3’. All genome edits were verified through Sanger sequencing.

### Statistical Analysis

All p-values were calculated using two-tailed t-tests assuming equal variance. All statistics are presented as mean ± one standard deviation.

### Data Availability Statement

All strains and reagents used in this study are available upon request. The authors affirm that all necessary data to support the conclusions of this study are included within the manuscript.

## Results

### Co-conversion of Tightly Linked Genes to Test Genome Editing Efficiency

In *C. elegans*, standard genome editing practices involve injection of Cas9, guide RNAs and homologous repair templates into the germline of self-fertile hermaphrodite adult animals (Xu, 2015; Chen et al., 2016; Dickinson and Goldstein, 2016; Farboud, 2017; Nance and Frokjaer-Jensen, 2019; Kim and Colaicovo, 2019; Iyer et al., 2018). Due to the syncytial nature of the distal gonad, a single injection can be distributed among numerous germ cells (Evans et al., 2006; Kadandale et al., 2008). Although injection of Cas9 into the distal germline is expected to affect the genomes of maternal oocytes, homozygously-edited animals can be recovered from the F1 generation post-injection, suggesting that editing of both maternal and paternal germ cells can occur from a single injection (Friedland et al., 2013; Zhao et al., 2014; Kim et al., 2014; Paix et al., 2015; Wang et al., 2018). In addition, PCR amplification of heterozygous animals may not amplify large deletions if the deletions affect primer binding sites (Katic and Grosshans 2013; Kim et al., 2014; Dokshin et al., 2018; Wang et al., 2018), further complicating quantification of genome editing rates by providing an inaccurate picture of editing nature (HDR or indel) and frequency. Overall, edited F1 animals can carry maternal genome edits, paternal genome edits, or both. Unless both maternal and paternal haplotypes can be analyzed separately, it can be difficult to determine whether one or two independent genome editing events may have occurred, complicating quantification of genome editing rates.

Therefore, we first aimed to develop a new method for quantifying genome editing rates that would allow separate analysis of each parental haplotype (Figure 1A). We used a co-editing (co-CRISPR) approach, using two tightly linked genes that were simultaneously targeted using two different guide RNAs (Figure 1A). An advantage of co-editing is that the editing of one locus ensures that Cas9 was active and available to target the second locus, which can at least partially normalize injection efficiencies across different injections (Kim et al., 2014; Paix et al., 2015). We chose to edit the *tra-2* gene, which is located on chromosome II, 0.16 map units away from the commonly used co-CRISPR gene *dpy-10*. The *dpy-10(cn64)* variation results in a semi-dominant phenotype that is easily visualized on a stereomicroscope (Arribere et al., 2014; Paix et al., 2015). Animals homozygous for the *dpy-10(cn64)* mutation have a dumpy phenotype marked by a reduced body length whereas heterozygous animals have a normal body length but display an abnormal rolling behavior (Figure 1A, Arribere et al., 2014; Paix et al., 2015).

**Figure 1.**
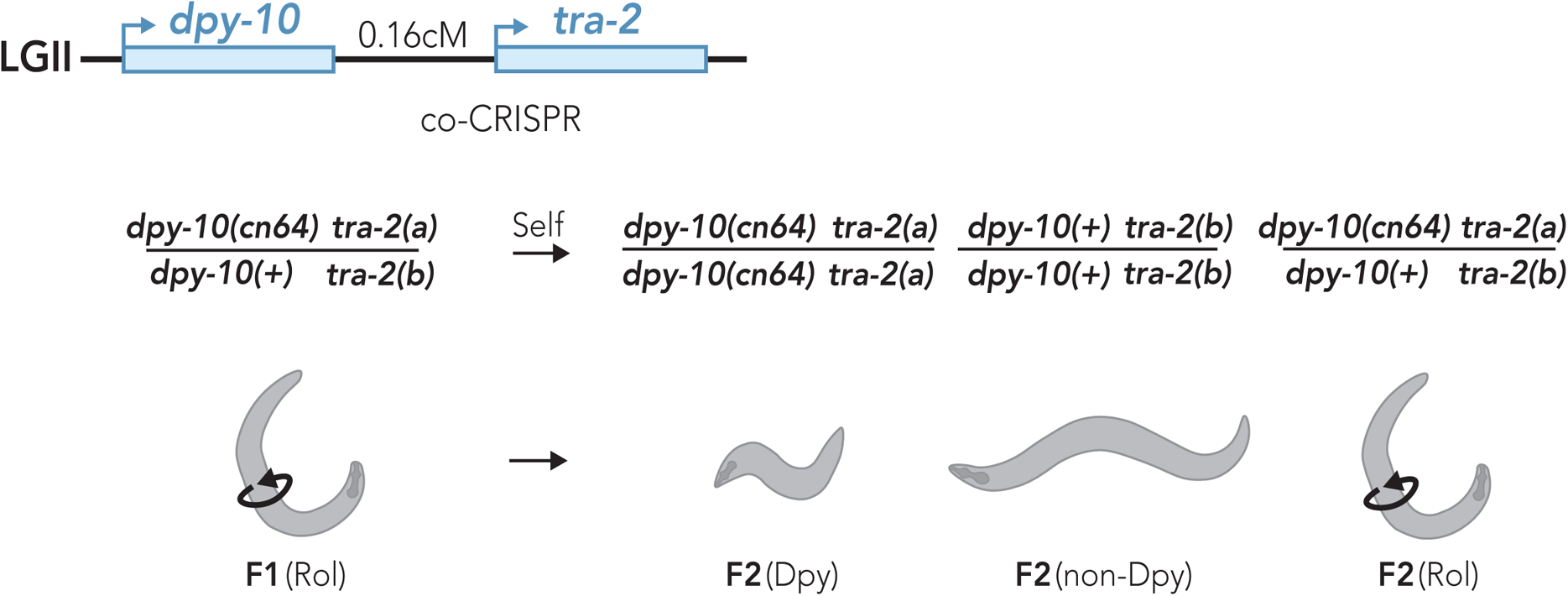
Strategy for co-conversion of *dpy-10* and *tra-2* to quantify haploid genome editing efficiency. *dpy-10* and *tra-2* are located 0.16 map units apart on LGII and do not undergo independent assortment (~1/625 meiotic recombination frequency). The *dpy-10(cn64)* allele produces a semi-dominant, physical phenotype where heterozygous animals have a rolling phenotype and homozygous animals have a dumpy phenotype. Following co-CRISPR of *dpy-10* and *tra-2*, one haplotype of F1 rollers has an unknown allele of *tra-2* (*‘tra-2(a)’*) linked to the *dpy-10(cn64)* variation and a second haplotype where an unknown *tra-2* allele (‘*tra-2(b)*’) is linked to a wildtype allele of *dpy-10*. Following self-fertilization of F1 hermaphrodite rollers, F2 generation dumpy animals are expected to be homozygous for the *tra-2(a)* allele whereas non-dumpy animals should be homozygous for the *tra-2(b)* allele.

Due to their close proximity, meiotic recombination between *tra-2* and *dpy-10* is only expected in 1/625 haplotypes. Therefore, the haplotype arrangement of *tra-2* and *dpy-10* alleles will be stably maintained across generations (Figure 1A). As F1 generation roller animals contain a single *dpy-10(cn64)*-marked (Dpy-marked) chromosome and one chromosome that is not Dpy-marked, we were able to distinguish the two parental haplotypes and determine whether either or both haplotypes carried an edited allele of *tra-2* (Figure 1A). Segregation of dumpy and non-dumpy animals in the F2 generation homozygoses for each F1 generation haplotype, which allowed us to definitively determine whether one or two genome editing events had taken place in the F1 generation by sequencing the *tra-2* genomic locus (Figure 1A).

### Single Nucleotide Blocking Mutations in the Guide-binding Region Allow for Effective Genome Editing

To determine whether single nucleotide substitutions in the guide-binding region effectively block Cas9 and allow for recovery of genome-edited animals, we designed a series of blocking mutations within a guide-binding region located in the 5’ UTR of *tra-2* (Figure 2A). Since previous reports have suggested that substitutions proximal to the 3’ end of the guide are more effective at blocking Cas9, (Jinek et al., 2012; Fu et al., 2013; Pattanayak et al., 2013; Hsu et al., 2013; Mali et al., 2013; Cong et al., 2013; Zhang et al., 2015), we introduced substitutions every three nucleotides to test the positional effects of single nucleotide blocking mutations (Figure 2A). We named each mutation according to its position (‘P’) relative to the 3’ end of the guide sequence (Figure 2A).

**Figure 2.**
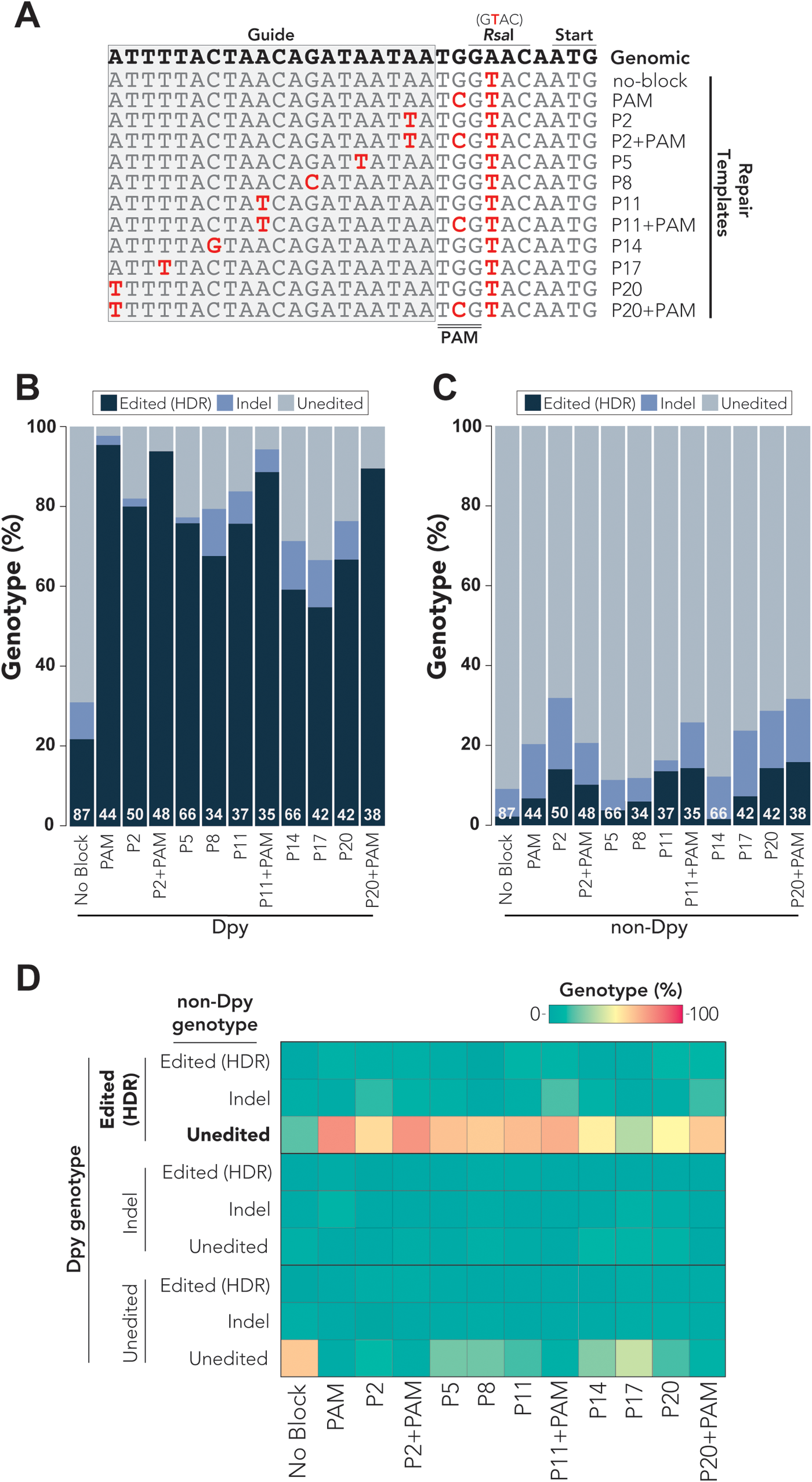
Single nucleotide mismatches in the guide RNA sequence are sufficient to promote homology-directed repair. (A) Partial alignment of *tra-2* repair templates used for genome editing experiments. See Table S2 for full length sequences of all repair templates. The wild-type genomic sequence is shown on the top line in bold text. Changes to the genomic sequence are indicated in red text. ‘P’ refers to position on the guide sequence (gray-shaded box) counting from the 3’ end of the guide. The PAM sequence is indicated by a double bar (below). (B-C) Percent *tra-2* genotypes observed for F2 generation dumpy (B) or non-dumpy (C) animals singled from F1 generation rollers. Edited (HDR) events were defined as partial or complete incorporation of genome edits engineered into single-stranded DNA repair templates. Indels were defined as any insertion or deletion mutation, regardless of whether editing through HDR may have also occurred. Unedited animals had no apparent changes compared to the wild-type *tra-2* genomic sequence. White text at the bottom of each stacked bar indicates the number (n) of animals that were sequenced. (D) Paired analysis of F2 generation dumpy and non-dumpy genotypes from a single F1 generation roller. Heatmap indicates the percent genotypes observed for each different blocking condition, where the sum of each column adds to 100% genotypes. Indel mutations were defined as any insertion or deletion mutation, regardless of whether editing through HDR may have also occurred. The most frequent observation (edited dumpy animals with unedited non-dumpy siblings) is bolded. All results were determined through Sanger sequencing of singled F2 generation dumpy or non-dumpy animals.

For example, we refer to the mutation affecting the second nucleotide from the 3’ end of the guide sequence as ‘P2’ and the mutation affecting the twentieth nucleotide from the 3’ end as ‘P20’ (Figure 2A). As a control, we designed a mutation within the PAM domain, which is expected to completely block Cas9 activity (Mojica et al., 2009; Marraffini et al., 2010; Sashital et al., 2012; Jinek et al., 2012). Each of the repair templates were designed to also include a non-blocking, single nucleotide substitution downstream of the PAM sequence that introduces an *Rsa*I restriction enzyme cutting site (Figure 2A). As an additional control, we designed a repair template that only included the *Rsa*I cutting site, which should not block Cas9 activity. This repair template would therefore not be expected to allow for HDR editing since Cas9 should continue targeting the repaired DNA.

We performed co-CRISPR of *dpy-10* and *tra-2* for each of the blocking conditions that we designed (Figure 2A). We used ssODN repair templates, which themselves are not subject to Cas9 cleavage and effectively promote genome editing when injected alongside pre-assembled Cas9 ribonucleotide protein (RNP) complexes in *C. elegans* (Paix et al., 2015). Following injection, we singled F1 generation roller animals from ‘jackpot’ broods containing the highest percentage of F1 generation *dpy-10*-edited animals (Paix et al., 2015). As rollers are heterozygous for the *dpy-10(cn64)* variation, F2 progeny are a mixture of homozygous dumpy animals, heterozygous roller animals and homozygous non-dumpy animals (Figure 1). Since *tra-2* and *dpy-10* are genetically linked, F2 generation dumpy and non-dumpy animals will each be homozygous for different alleles of *tra-2* (Figure 1). We then sequenced the *tra-2* genomic locus of the F2 animals to determine whether a single haplotype or both haplotypes had been edited. We scored the frequency of HDR editing and the frequencies of insertion or deletion (indel) mutations (Figure 2, B-C, Table S3). We also quantified the number of animals that did not show any apparent editing of *tra-2* (unedited) (Figure 2, B-C, Table S3). We defined HDR-edited animals as any animal that incorporated any of the designed mutations, regardless of whether partial or complete repair had occurred. As the *tra-2* guide RNA targets the 5’ UTR of *tra-2*, relatively small indel mutations might lead to loss of *tra-2* function. Loss of *tra-2* is not lethal but results in masculinization of hermaphrodite animals (Hodgkin and Brenner, 1977; Doniach, 1986), which allowed us to recover deleterious mutations such as indels.

In F2 generation dumpy animals, we observed nearly complete HDR editing of *tra-2* when the PAM was mutated (95.4% HDR-edited, Figure 2B, Table S3), which is consistent with PAM mutations blocking Cas9 activity (Mojica et al., 2009; Marraffini et al., 2010; Sashital et al., 2012; Jinek et al., 2012). We observed similarly high HDR-editing rates in F2 generation dumpy animals when the PAM was mutated alongside an additional single nucleotide mismatch in the guide-binding region such as P2+PAM (93.8% HDR-edited), P11+PAM (88.6% HDR-edited) or P20+PAM (89.5% HDR-edited, Figure 2B, Table S3). Although the efficiency was slightly reduced compared to the PAM blocking conditions, we found that the P2 (80.0% HDR-edited), P5 (75.8% HDR-edited), P8 (67.6% HDR-edited), P11 (75.7% HDR-edited), P14 (59.1% HDR-edited), P17 (54.8% HDR-edited) and P20 (66.6% HDR-edited) single nucleotide guide substitutions allowed for effective HDR editing in F2 generation dumpy animals (Figure 2B, Table S3). The position of the single nucleotide substitutions within the guide-binding region appears to influence their efficacy at promoting HDR, as we found that substitutions closer to the 3’ end of the guide sequence tended to be more effective at promoting HDR compared to substitutions near the 5’ end of the guide sequence (Figure 2B, Table S3). For example, the P2 (80.0% HDR-edited), P5 (75.8% HDR-edited), and P8 (67.6% HDR-edited) substitutions located within the 3’ half of the guide sequence averaged significantly higher HDR-editing rates (74.5 ± 6.3% average HDR-edited) compared to the P14 (59.1% HDR-edited), P17 (54.8% HDR-edited) and P20 (66.6% HDR-edited) substitutions that are located in the 5’ half of the guide sequence (60.2% average HDR-edited, p<0.05). This positional effect of guide substitutions was not surprising given that previous studies have demonstrated that mismatches near the 3’ end of the guide sequence are more effective at blocking Cas9 than mismatches near the 5’ end of the guide sequence (Jinek et al., 2012; Fu et al., 2013; Pattanayak et al., 2013; Hsu et al., 2013; Mali et al., 2013; Cong et al., 2013; Zhang et al., 2015).

Interestingly, for all blocking conditions examined, we noticed a substantial reduction in HDR editing efficiency of the *tra-2* locus in F2 generation non-dumpy animals (Figure 2C, Table S3) compared to their dumpy siblings (Figure 2B, Table S3). For example, whereas the PAM blocking condition resulted in very high HDR-editing rates in F2 generation dumpy animals (95.4% HDR-edited, Figure 2B, Table S3), the rate of HDR editing was much lower in their non-dumpy siblings (6.8% HDR-edited, Figure 2C, Table S3). Although all single nucleotide substitutions in the guide-binding region had lower HDR editing rates in F2 generation non-dumpy animals compared to dumpy animals, we did not observe a strong correlation between the position of substitutions within the guide sequence and the efficiency of HDR editing for non-dumpy animals (Figure 2C, Table S3), which is in contrast to the positional effects that we observed in their dumpy siblings (Figure 2B, Table S3). While the substitutions in the 3’ half of the guide sequence (P2, P5 and P8) were more effective at promoting HDR than substitutions in the 5’ half of the guide sequence (P14, P17 and P20) in dumpy animals (Figure 2B, Table S3), the same substitutions in the 3’ half of the guide sequence resulted in a similar HDR-editing rate (8.67 ± 4.62% average HDR-edited) as substitutions in the 5’ half of the guide sequence (7.67 ± 6.41% average HDR-edited, p=0.83) for the F2 generation non-dumpy siblings (Figure 2C, Table S3). Importantly, we were able to recover HDR-edited animals for all blocking mutations that we tested in F2 generation dumpy and non-dumpy animals, showing that single nucleotide guide substitutions are sufficient to allow for effective HDR-editing in *C. elegans* and can be used as an alternative to PAM mutations when silently mutating the PAM is not possible.

We also quantified the frequency of indel mutations that occurred in the *tra-2* locus for each blocking condition that we tested (Figure 2, B-C, Table S3). Since indels often result from NHEJ repair pathways, the presence of indels might suggest that NHEJ had been favored over HDR, which might be expected under conditions where Cas9 was not completely blocked. Consistent with this idea, we observed low indel rates in F2 generation dumpy animals when the PAM was mutated (2.3% indels) or when the PAM was mutated alongside an additional mutation in the guide sequence: P2+PAM (0.0% indels), P11+PAM (5.7% indels) or P20+PAM (0.0% indels, Figure 2B, Table S3). By comparison, we observed slightly increased indel rates when using single nucleotide blocking mutations within the guide sequence in F2 generation dumpy animals (8.1 ± 4.6% indels, p<0.05, Figure 2B, Table S3). Furthermore, the position of the substitutions within the guide sequence appears to influence the frequency of indels observed under each blocking condition (Figure 2B, Table S3). For example, we observed a low indel rate for the P2 (2.0% indels) and P5 (1.5% indels) substitutions, located near the 3’ end of the guide sequence, whereas we observed increased indel frequency P17 (11.8% indels) and P20 (9.6% indels) substitutions that are closest to the 5’ end of the guide sequence (Figure 2B, Table S3). Thus, the expected blocking efficiency of each mutation appears to inversely correlate with the frequency of indel mutations under that blocking condition. In most cases, the indel rate in F2 generation non-dumpy animals was increased compared to the respective blocking mutations in their dumpy siblings (Figure 2, Table S3) For example, while the P2 mutation led to a 2.0% indel rate in dumpy animals, this rate increased to 18.0% in non-dumpy animals (Figure 2, Table S3). This suggests that non-dumpy animals are biased towards NHEJ-dependent repair pathways compared to their dumpy siblings. This observation suggests that different mechanisms might influence repair of the *dpy-10*-edited haplotypes compared to the haplotypes that are not edited at the *dpy-10* locus.

We were surprised to find that we were still able to recover HDR-edited animals when a blocking mutation was not incorporated into the repair template. We found that 21.8% of dumpy and 2.3% of non-dumpy F2 generation animals carried HDR-edited mutations (Figure 2, Table S3), with HDR-editing assessed by the presence of the non-blocking *Rsa*I restriction site. Although the HDR-editing rates under the no-blocking condition were substantially reduced compared to conditions that introduced a blocking mutation (Figure 2, Table S3), it was unexpected that any HDR-edited animals could be recovered given that Cas9 was not blocked from targeting the repaired DNA. Thus, blocking mutations do not appear to be absolutely required to promote genome editing via HDR in *C. elegans*, suggesting that a temporal block might restrict Cas9 from continuing to target the genome after HDR.

We next examined whether there was a correlation between the *tra-2* genotypes of dumpy and non-dumpy F2 generation sibling animals that originated from the same F1 hermaphrodite animal. Toward this end, we performed a paired analysis of the *tra-2* genotypes that we observed in F2 generation animals (Figure 2D). Under all blocking conditions that we examined, the majority of F2 generation dumpy animals were HDR-edited, whereas their non-dumpy siblings were usually non-edited (Figure 2D, Table S4) In other words, the non-dumpy sibling had no apparent changes to the wild-type *tra-2* genomic sequence. Interestingly, when the F2 generation dumpy animal had an indel mutation the non-dumpy sibling did not appear to be more likely to have an indel mutation, suggesting that the editing of each *tra-2* haplotype occurs independently (Figure 2D, Table S4). For example, under the P2 blocking condition the indel rates of non-dumpy animals were highest when HDR editing had occurred in their dumpy sisters (14.3% indels) but were lower when the dumpy sister had an indel mutation (0.0% indels) or were not edited (5.4% indels) (Figure 2D, Table S4). Conversely, in the PAM blocking condition the indel rates of non-dumpy animals were highest when their dumpy sister also had an indel mutation (9.3% indels) but were lower when their dumpy sister was HDR-edited (2.3% indels) or not edited (0.0% indels) (Figure 2D, Table S4). When no blocking mutation was introduced into the repair template, we found that the most frequently observed combination of genotypes was that both dumpy and non-dumpy sisters were not edited (Figure 2D, Table S4). Thus, editing of each *tra-2* haplotype, one each of maternal and paternal origin, appears to occur independently.

### The Position of Single Nucleotide Blocking Mutations Influences the Completeness of Homology-Directed Repair

Although we observed high HDR-editing rates among all of the blocking conditions that we tested, we found that partial HDR-dependent repair often occurred, where only a subset of the designed mutations was incorporated into the genome (Figure 3A, Table S5). For example, in some cases, the blocking mutation was incorporated while the *Rsa*I restriction site had not been edited and vice versa (Figure 3A, Table S5). We next asked whether the position of blocking mutations influenced the efficacy of HDR (Figure 3A, Table S5). For the purpose of this analysis, we only considered genotypes where partial or complete HDR-editing had occurred (Figure 3A, Table S5). We quantified the percentage of HDR-edited genotypes that contained a designed blocking mutation, the *Rsa*I restriction site, or both blocking and *Rsa*I mutations (Figure 3A, Table S5). We found that blocking conditions where the PAM was mutated led to increased incorporation of both the blocking mutation and *Rsa*I restriction site among the HDR-edited animals (Figure 3A, Table S5). When only the PAM was mutated, the majority of HDR-edited chromosomes contained both the PAM blocking mutation and the *Rsa*I restriction site (97.8% both mutations), while a small percentage contained only the *Rsa*I restriction site (2.2% *Rsa*I only) (Figure 3A, Table S5). Interestingly, all HDR-edited chromosomes that we examined for the P2+PAM blocking condition were edited for both the blocking mutation and the *Rsa*I restriction site (100% both mutations), while the P2 blocking condition itself resulted in less frequent incorporation of both blocking mutations and the *Rsa*I restriction site (36.2% both mutations) and commonly resulted in partial repair of only the blocking mutation (31.9% P2 only) or the *Rsa*I restriction site (31.9% *Rsa*I only) (Figure 3A, Table S5). We observed a similar trend for the P11+PAM or P20+PAM blocking conditions compared to the P11 or P20 blocking conditions respectively, where complete HDR of both blocking mutations and the *Rsa*I restriction site was more frequent when the PAM was also mutated (Figure 3A, Table S5). While we cannot be certain why the presence of PAM blocking mutations led to increased frequency of complete HDR repair compared to blocking mutations in the guide sequence, we hypothesize that partial HDR repair could reflect incomplete blocking of Cas9. This, in turn, could lead to Cas9 recutting of repaired DNA and increased frequency of inaccurate HDR as repair is attempted multiple times. We also observed that HDR appeared to occur in an asymmetric fashion, favoring blocking mutations over *Rsa*I site incorporation. For example, when P20 was used as a blocking mutation we observed more frequent incorporation of the P20 mutation (87.5% P20 edited) than the *Rsa*I restriction site (46.9% *Rsa*I edited), despite *Rsa*I being located closer to the expected Cas9-generated break site (Figure 3A, Table S5). This observation suggests that edits that do not block Cas9 from recutting (such as introduction of the *Rsa*I site) may result in multiple cleavage and repairs, thus ultimately favoring repair events that incorporate the blocking mutations. However, the presence of *Rsa*I edits that do not include blocking mutations further supports the existence of a temporal block that prevents Cas9 from recutting the repaired template.

**Figure 3.**
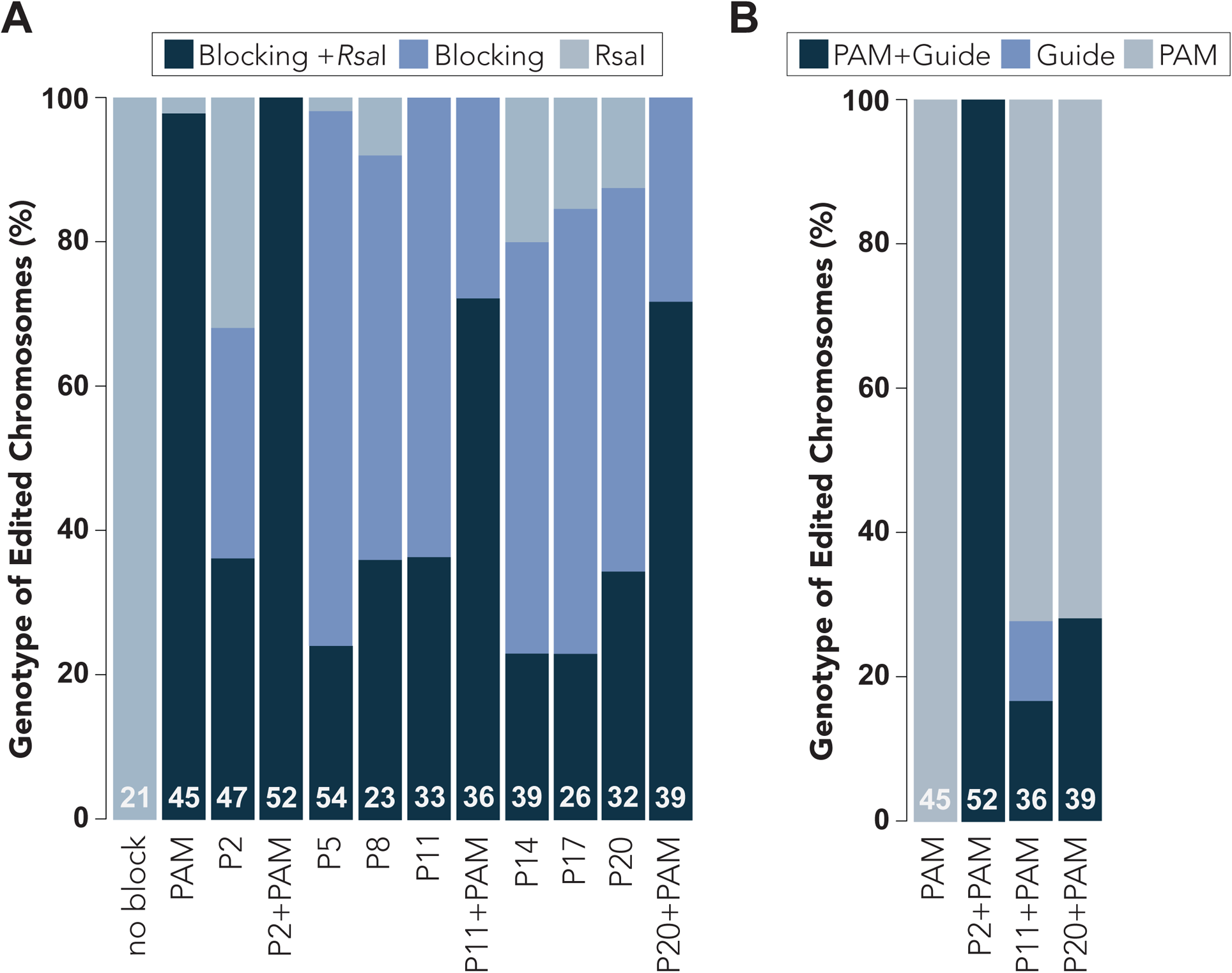
Differences in HDR Editing Efficiency Under Different Blocking Conditions. (A) Percent HDR-edited chromosomes containing mutations for the non-blocking *Rsa*I restriction enzyme cutting site, blocking mutation, or both blocking and *Rsa*I mutations. For genome edits that introduce more than one blocking mutation into the PAM and guide sequence, we scored the presence of at least one blocking mutation (i.e. PAM, guide, or both). (B) Percent HDR edited chromosomes containing blocking mutations in the PAM sequence, guide sequence, or both PAM and guide sequences. (A-B) All results were determined through Sanger sequencing of singled F2 generation animals. White text at the bottom of each stacked bar indicates the number (n) of animals that were sequenced.

Since previous reports have shown that the efficiency of HDR editing is inversely correlated with distance to the Cas9-generated break site (Inui et al., 2014; Arribere et al., 2014; Paix et al., 2014; Ward, 2015; Paix et al., 2015), we asked whether the distance of blocking mutations from the break site might be confounded with their actual ability to block Cas9. In other words, the P2 blocking mutation might be more effective at promoting HDR than the P20 blocking mutation because it is closer to the dsDNA break site and is therefore more likely to incorporate during repair, rather than the P2 mutation blocking Cas9 more effectively than P20. To address this, we examined how frequently partial or complete HDR occurred when using repair templates that mutated one nucleotide within the guide sequence as well as the PAM domain to ensure that Cas9 was equally blocked under each condition (P2+PAM, P11+PAM and P20+PAM) (Figure 3B, Table S6). When both the PAM and P2 mutations were introduced, we found that all of the edited chromosomes that we examined contained both PAM and P2 mutations (100% both edited) (Figure 3B, Table S6). However, when the PAM was mutated alongside the P11 or P20 mutations, we found that partial HDR repair had often occurred. In particular, we found that the majority of HDR-edited chromosomes contained only the PAM blocking mutation when either P11+PAM (72.2% PAM only) or the P20+PAM (71.8% PAM only) repair templates had been used (Figure 3B, Table S6). Only a small fraction of HDR-edited chromosomes contained both the guide blocking mutation and the PAM mutation for the P11+PAM (16.6% both edited) or P20+PAM (28.2% both edited) mutations (Figure 3B, Table S6). Therefore, the distance of blocking mutations from the dsDNA break site, which is located near the 3’ end of the guide sequence, appears to influence the rate at which they are incorporated via HDR. Blocking mutations located farther away from the dsDNA break site may not always be incorporated through HDR, leading to an overall decrease in HDR-editing efficiency since Cas9 can re-target the partially repaired chromosome. Despite the reduced incorporation of the more distant blocking mutations, the overall high efficiency of obtaining desired edits (Figure 2) strongly supports practical use of single nucleotide blocking mutations when PAM mutations are not possible.

### Single Nucleotide Guide Substitutions Effectively, Although not Completely, Block Cas9

Having determined that single nucleotide blocking mutations can promote HDR (Figure 2), we next asked how effective each of blocking mutations might be in preventing Cas9 from targeting the *tra-2* genomic locus after HDR occurred. To address this question, we took advantage of the *tra-2* HDR-edited animals that we generated (Figure 2), containing blocking mutations as well as the *Rsa*I restriction enzyme cutting site. We reasoned that if the blocking mutations built into the *tra-2* locus prevent Cas9 from targeting the mutated sequences, then re-injection of Cas9 and the same guide RNA, perfectly matched to the wild-type *tra-2* genomic sequence, should not lead to editing of the mutated *tra-2* loci (Figure 5A). To examine whether re-editing of *tra-2* could occur under any of the blocking conditions, we designed a repair template that would revert *tra-2* back to the wild-type genomic sequence (Figure 4A). Since the HDR-edited strains contain blocking mutations and the *Rsa*I restriction enzyme site, reversion of *tra-2* back to the wild-type sequence will result in removal of the *Rsa*I restriction enzyme site (Figure 4A). Furthermore, as the *Rsa*I restriction enzyme site is located immediately downstream of the *tra-2* PAM (Figure 2A), indel mutations might also be expected to disrupt the *Rsa*I site. On the other hand, if Cas9 is completely blocked from generating a dsDNA break at the mutated *tra-2* locus, then all of the F1 progeny post-injection should still contain the *Rsa*I restriction enzyme site. Importantly, any animal now lacking the *Rsa*I site must have been targeted by Cas9 for genome editing to have occurred, which would indicate that Cas9 was not completely blocked under that condition. To test whether the position of blocking mutations within the guide sequence influenced their blocking efficacy, we examined the *Rsa*I reversion efficiencies of the P2, P11 and P20 mutations (Figure 4A). As a positive control, we reverted the *Rsa*I site in a strain that did not contain any Cas9 blocking mutations (‘no-block’), which would still be expected to be targeted by Cas9 loaded with the wild-type guide (Figure 4A). Since the PAM is absolutely required for Cas9 activity (Mojica et al., 2009; Marraffini et al., 2010; Sashital et al., 2012; Jinek et al., 2012), we attempted to revert the *Rsa*I site in PAM-edited animals as a negative control (Figure 4A).

**Figure 4.**
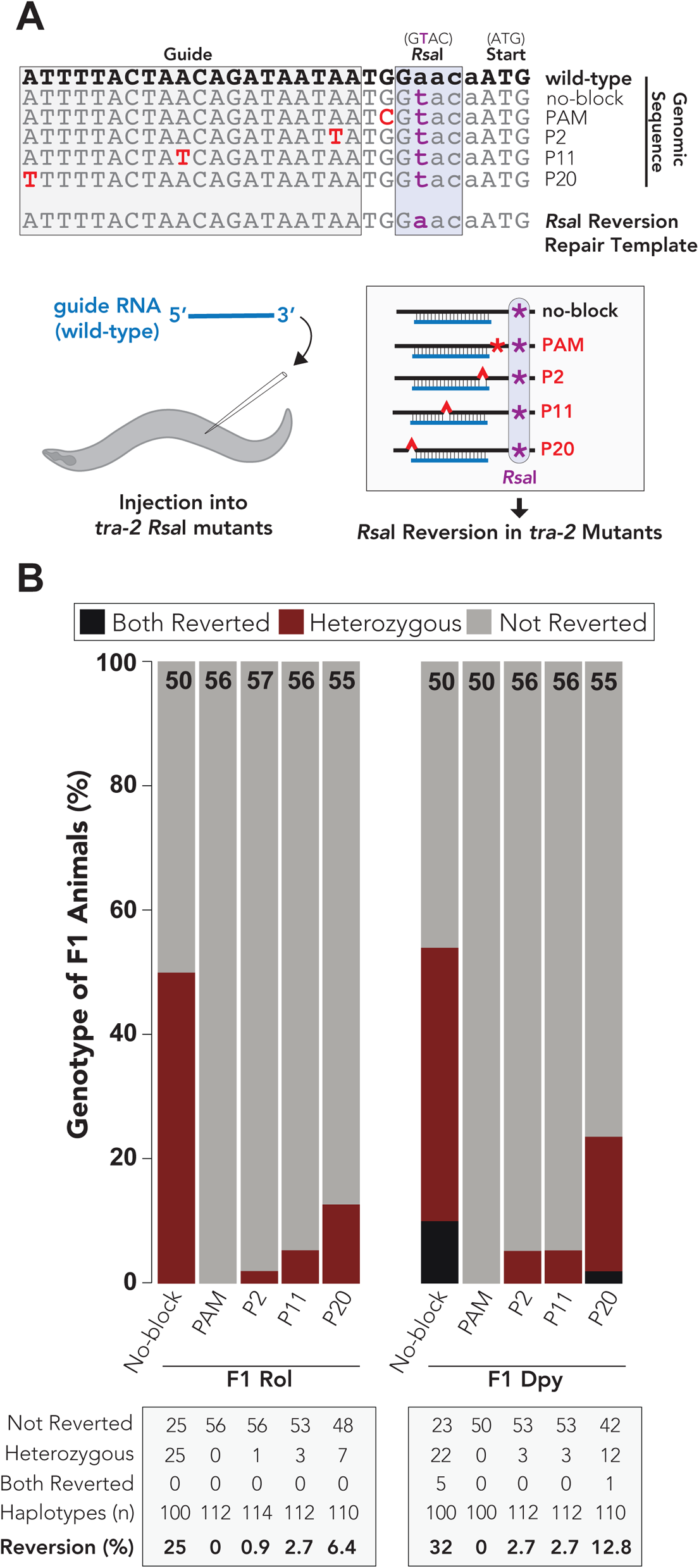
Single nucleotide guide substitutions effectively block Cas9 in a position-dependent manner. (A) Top: sequence alignment of *tra-2* genome edited animals that were re-injected to revert the *tra-2 Rsa*I restriction site back to the wild-type genomic sequence. Note that the same *Rsa*I reversion repair template was injected into all strains and is also expected to revert the blocking mutations back to the wildtype genomic sequence. As the *Rsa*I cutting site is located nearby the expected cutting site for Cas9, both *Rsa*I reversions and relatively small deletions would be expected to eliminate *Rsa*I cutting for edited alleles. Since some mutations would not disrupt *Rsa*I digestion, this analysis underestimates the true mutation rate. Bottom: schematic representation of the experimental procedure. Animals containing tra-2 RsaI mutations are injected with a wild-type guide. As the tra-2 mutants are genome edited, the wild-type guide sequence is not perfectly paired to the mutant genomic sequences (B) Quantification of F1 generation roller and dumpy animals digested with *Rsa*I. “Both reverted” indicates that the singled F1 animals were homozygous for *Rsa*I reversion back to wild type (did not digest with *Rsa*I). “Heterozygous” animals showed both patterns of *Rsa*I digestion. “Not reverted” indicates that the animals underwent full *Rsa*I digestion and were not reverted back to wild type. Black text at the top of each stacked bar indicates the number (n) of F1 animals that were digested with *Rsa*I. Table (bottom) shows number of F1 generation animals corresponding to each genotype. Reversion (%) illustrates the haplotype editing frequency, which was calculated by dividing the total number of edited haplotypes (two in ‘both reverted’ and one in heterozygous animals) from the total number of haplotypes examined.

**Figure 5.**
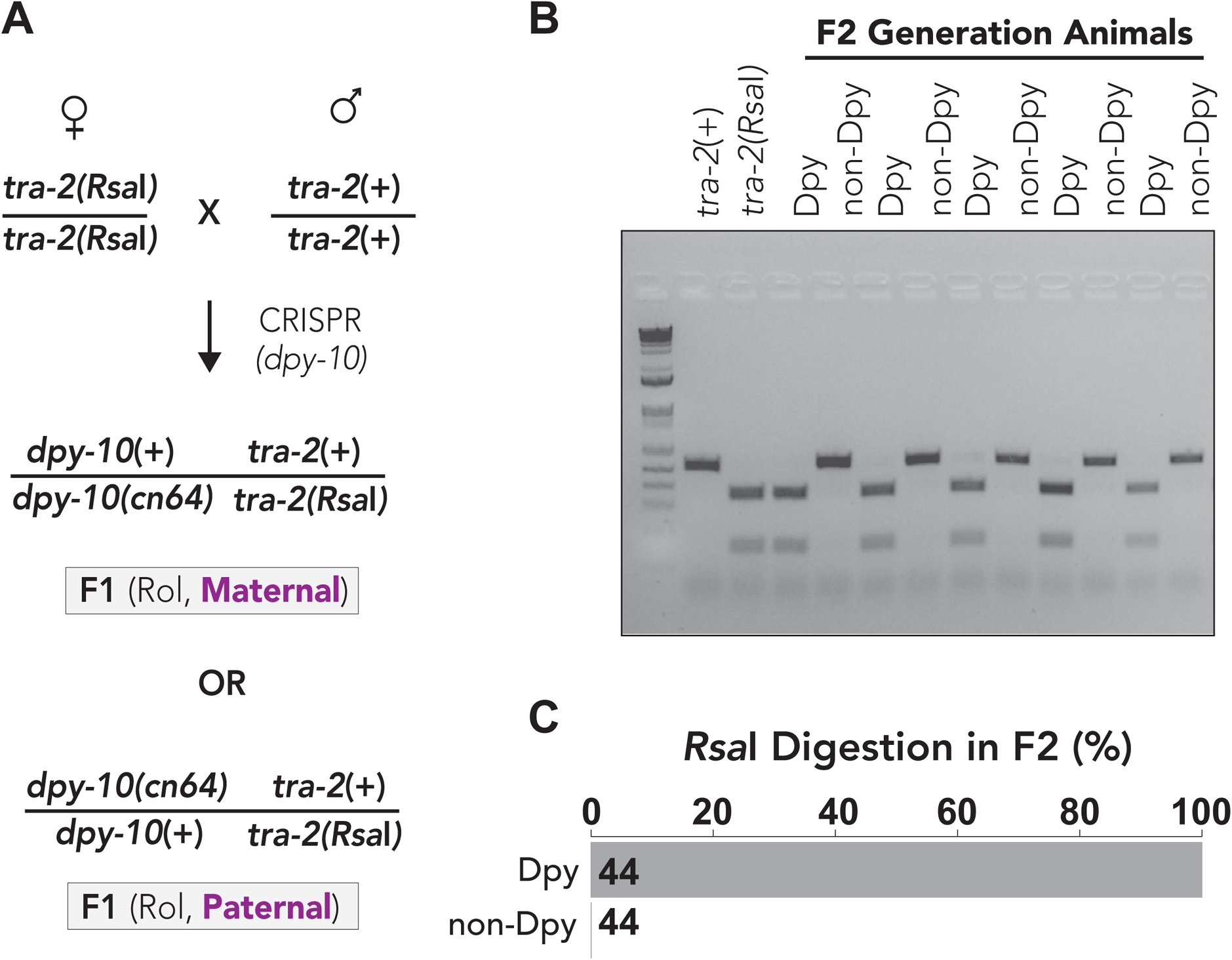
Analysis of maternal and paternal genome editing rates. (A) Mating strategy to test frequency of maternal versus paternal genome editing. Hermaphrodite animals containing the *tra-2 Rsa*I (no-blocking) restriction enzyme site were crossed to wild-type males and allowed 24h to mate. Mated hermaphrodites were subsequently injected to generate the *dpy-10(cn64)* variation. F1 generation rollers contain a single edit for the *dpy-10(cn64)* allele and a maternally provided allele of *tra-2* that contains the *Rsa*I site. For maternal edits, the F2 generation dumpy animals will be homozygous for the *tra-2* allele containing the *Rsa*I restriction site. In the case of paternal edits, the non-dumpy animals will be digested with *Rsa*I. (B) Representative agarose gel illustrates differences in editing efficiencies of maternal and paternal genomes. Wildtype animals are not digested with *Rsa*I and migrates as a single band whereas animals containing the *tra-2(Rsa*I) allele are digested and migrate as two bands. (C) Quantification of *Rsa*I digestion rates in F2 generation dumpy and non-dumpy animals. *Rsa*I digestion in dumpy animals is indicative of maternal genome editing of *dpy-10*. Black number indicates the number (n) of animals that were digested with *Rsa*I.

We used *dpy-10* co-conversion to enrich for genome-edited animals and examined whether F1 generation roller and dumpy animals contained the *tra-2 Rsa*I restriction site (Figure 4B, Table S7). We calculated the haplotype *Rsa*I reversion rate by dividing the number of reverted haplotypes (one for heterozygous reverted animals and two for homozygous reverted animals) by the total number of haplotypes that were examined.

As expected, we observed robust reversion of the *Rsa*I restriction site in the no-blocking control and never observed reversion of the *Rsa*I site in PAM-edited animals (Figure 4B). For the no-blocking control, we found that the *Rsa*I reversion rates of F1 generation dumpy (32% haplotypes reverted) and roller (25% haplotypes reverted) animals were similar, although we only observed homozygous-reverted animals in F1 generation dumpy animals (Figure 4B, Table S7). The increased prevalence of homozygous-reverted animals in F1 generation dumpy animals might not be surprising, since the dumpy phenotype indicates that homozygous editing of the *dpy-10* genomic locus had also occurred. We found that the P2 blocking condition was highly effective at blocking Cas9, as only a small percentage of F1 generation dumpy (2.7% haplotypes reverted) or roller (0.9% haplotypes reverted) animals had reverted the *Rsa*I site (Figure 4B, Table S7). Similarly, the P11 mutation was highly effective at blocking Cas9 and resulted in only a low frequency of *Rsa*I reversion in F1 generation dumpy (2.7% haplotypes reverted) and roller (2.7% haplotypes reverted) animals (Figure 4B, Table S7). Despite increased *Rsa*I reversion frequency in P20 blocking mutants for both F1 generation dumpy (12.8% haplotypes reverted) and roller (6.4% haplotypes reverted) animals (Figure 4B, Table S7), the P20 blocking mutation blocked Cas9, as the *Rsa*I reversion rates were much lower in P20 mutants compared to no-block controls (Figure 4B, Table S7). Therefore, consistent with previous reports (Jinek et al., 2012; Fu et al., 2013; Pattanayak et al., 2013; Hsu et al., 2013; Mali et al., 2013; Cong et al., 2013; Zhang et al., 2015), the position of single nucleotide guide substitutions appears to influence their blocking efficacy where substitutions located proximal to the 3’ end of the guide are more effective at blocking Cas9 (Figure 4B, Table S7). Interestingly, although we observed similar HDR editing rates for the P11 (75.7% HDR-edited in Dumpy animals, Figure 2, Table S3) and P20 mutations (66.6% HDR-edited in Dumpy animals, Figure 2, Table S3), P11 was more effective at blocking Cas9 than P20 (Figure 4B, Table S7). This discrepancy between blocking efficacy and HDR editing rates further supports the idea that perhaps a temporal effect contributes toward allowing for robust genome editing when Cas9 is not completely blocked. Collectively, these findings show that single nucleotide substitutions in the guide sequence effectively, albeit not completely, block Cas9 after HDR occurs, and that conditions where Cas9 is not completely blocked still allow for efficient genome editing in *C. elegans*.

### Editing of Maternal Haplotypes Occurs at Greater Frequency than Editing of Paternal Haplotypes

Given that we observed significantly higher HDR-editing rates of the *tra-2* locus in *dpy-10*-edited haplotypes compared to non-*dpy-10* haplotypes (Figure 2B-D), we asked whether there could be differences in the HDR-editing efficiencies of each parentally contributed haplotype. To determine the HDR editing rates of each parental haplotype, we used a mating-based approach to differentiate between maternal and paternal haplotypes (Figure 5A). We crossed wildtype males to *tra-2* mutant hermaphrodites that carried the *Rsa*I restriction site just upstream of the *tra-2* coding sequence (Figure 4A). We then injected the mated animals with Cas9 RNP complex to generate the *dpy-10(cn64)* variation (Figure 5A). Since *dpy-10* and *tra-2* are genetically linked and not expected to independently assort, the resulting F1 generation roller animals will have two possible haplotype arrangements for the *dpy-10* and *tra-2* mutations (Figure 5A).

One possible F1 genotype will have the maternally contributed *tra-2* mutation (*Rsa*I site) on the same chromosome as the newly introduced *dpy-10(cn64)* mutation and another possible F1 genotype would have the *tra-2* and *dpy-10* mutations on different chromosomes (Figure 5A). Both of these possible haplotype arrangements can be differentiated in the F2 generation by examining whether the dumpy or non-dumpy progeny contain the *tra-2 Rsa*I restriction site (Figure 5A). If maternal editing of *dpy-10* occurs, then the F2 generation dumpy animals would be homozygous for the *tra-2 Rsa*I allele whereas the non-dumpy F2 generation animals would be homozygous for the paternally provided wild type *tra-2* allele (Figure 5A). If editing occurred in the paternal genome, then the F2 generation dumpy animals would be homozygous for the wild type *tra-2* allele that lacks the *Rsa*I cut site (Figure 5A). Strikingly, in all F2 generation animals (n=44 animals) that we examined, *Rsa*I digestion occurred only in dumpy animals, supporting the idea that editing of the maternal genome is strongly preferred over paternal genome editing (Figure 5, B-C). Thus, we conclude that the distinct *tra-2* HDR rates we previously observed in dumpy versus non-dumpy F2 generation animals (Figure 2, B-C) are likely due to differences in the editing efficiencies of each paternal genome, where dumpy animals are likely homozygous for the maternally provided haplotype and non-dumpy animals are likely homozygous for the paternally provided haplotype.

### Selection of maternally provided haplotypes using balancer chromosomes to mutate the *let-7* miRNA in a scarless fashion

As we observed increased HDR editing efficiency of maternally provided genomes compared to paternally provided genomes (Figure 5), we devised a strategy to select for the maternally provided haplotypes post-injection (Figure 6A). Our approach was to label paternally provided chromosomes prior to injection by mating wild type hermaphrodite animals to males expressing fluorescently labelled balancer chromosomes (Figure 6A). Importantly, the use of balancer chromosomes restricts recombination between the maternally provided wildtype chromosome and the paternally provided balancer chromosome. This allows for segregation of each parental haplotype in subsequent generations and easily homozygoses for the desired edit, thereby reducing screening efforts post-injection (Figure 6A). We used this strategy, along with single nucleotide blocking mutations to mutate the *let-7* miRNA in a scarless fashion. As loss of *let-7* function is lethal (Reinhart et al., 2000), this strategy also allowed us to immediately maintain deleterious *let-7* mutations in a balanced, heterozygous genetic background, using the *tmc24* balancer chromosome (Dejima et al., 2018). Since the *tmc24* balancer contains a *pmyo-2::Venus* fluorescent marker that is pharyngeal expressed (Dejima et al., 2018), F1 generation cross-progeny should be fluorescently labelled whereas self-progeny would not be labelled (Figure 6A). In the subsequent F2 generation, animals without expression of pharyngeal Venus should be homozygous for the maternally provided *let-7* haplotype as the non-Venus animals lack the paternally provided *tmc24* chromosome (Figure 6A).

**Figure 6.**
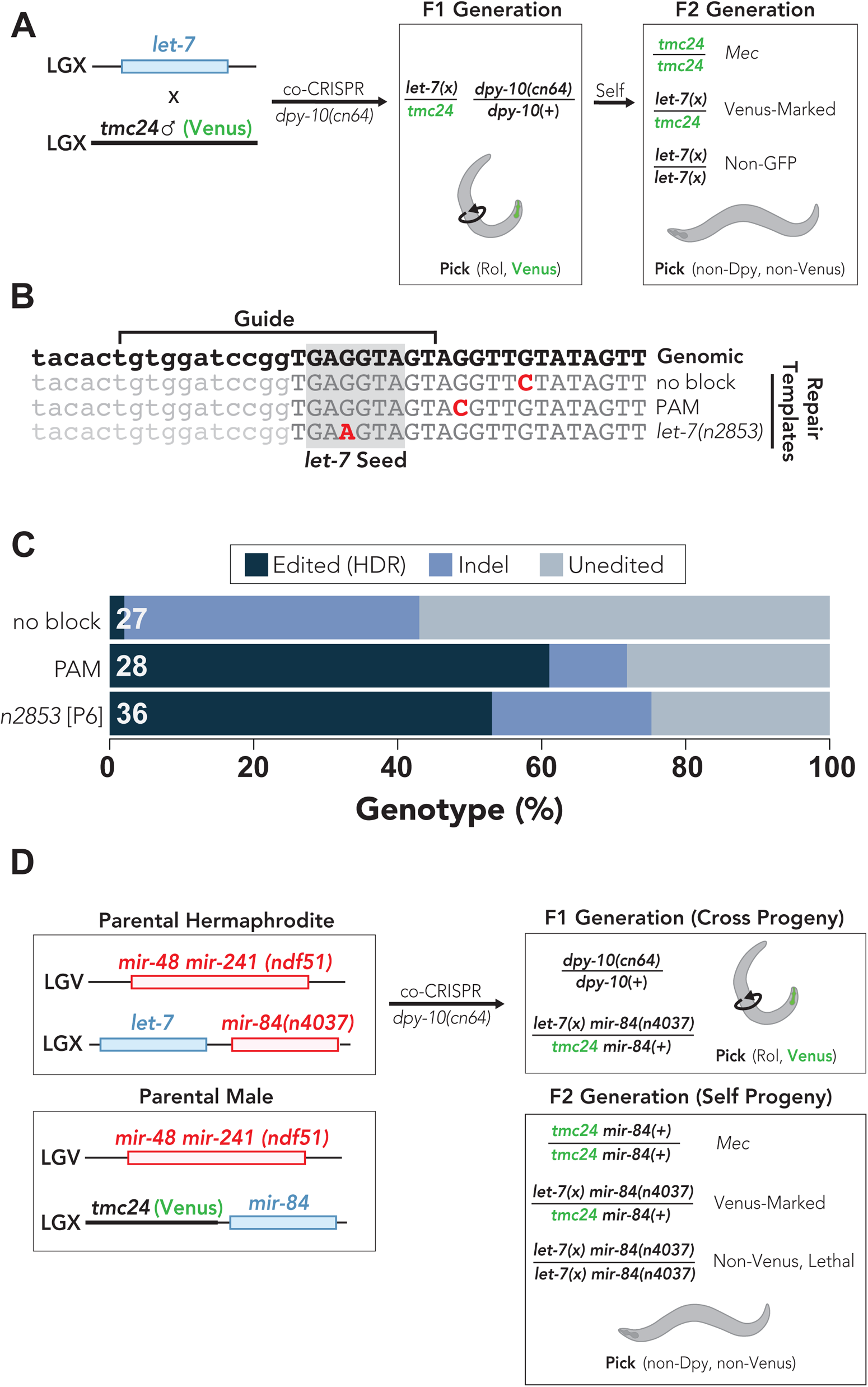
Selection of maternally provided haplotypes using balancer chromosomes. (A) Mating wildtype hermaphrodites to males carrying balancer chromosomes before injection allows for differentiation of maternal and paternal haplotypes for genes that are within the balanced interval and takes advantage of the more frequent maternal edits. The *tmc24* balancer covers an interval on the right side of LGX that includes the *let-7* miRNA and contains a Venus-marked transgene expressed in the pharynx. Prior to injection, *tmc24* males were mated to wild-type hermaphrodites and then subjected to co-CRISPR to mutate the *let-7* and *dpy-10* genes. F1 generation cross progeny will contain pharyngeal Venus, and roller animals were successfully mutated in the *dpy-10* gene. Since *dpy-10* and *let-7* are not on the same chromosome, non-dumpy animals can be isolated in the F2 generation to remove the *dpy-10(cn64)* allele. Homozygous animals for the *tmc24* balancer have a mechanosensory variant (Mec) phenotype, which can differentiate homozygous and heterozygous animals. Non-Venus F2 generation animals do not contain the paternally contributed *tmc24* balancer and are therefore homozygous for the maternally contributed *let-7* allele. (B) Alignment of *let-7* genome edits generated in this study. The wild-type genomic sequence is shown on the top line. The mature *let-7* miRNA is indicated by uppercase lettering and the *let-7* seed sequence is boxed in gray. Changes to the genomic sequence are indicated in red text. Introduction of the *let-7* seed mutation equivalent to the *let-7(n2853)* allele, which leads to a single nucleotide variation 6 bases away from the 3’ end of the guide sequence (‘P6’). (C) Percent *let-7* genotypes observed for F2 generation non-Venus animals that were singled from F1 generation Venus-positive rollers. Indels were defined as any insertion or deletion mutation, regardless of whether editing through HDR may have occurred. Unedited animals had no apparent changes compared to the wild-type *let-7* sequence. All results were determined through Sanger sequencing. White text at the left of each stacked bar indicates the number (n) of animals that were sequenced. (D) Strategy to generate *let-7* family *mir-48 mir-241(nDf51); mir-84(n4037) let-7(n2853)-*equivalent quadruple mutant. Hermaphrodite animals containing *mir-48 mir-241* and *mir-84* deletions are crossed to *ndf51; tmc24* males prior to injection, which homozygoses *ndf51* in subsequent generations. Note that mir-84 is on chromosome X and is genetically linked to the tmc24 balancer. While *mir-84* is not on the balanced interval covered by *tmc24*, *mir-84* and *tmc24* are not expected to independently assort during meiosis, which effectively maintains the *mir-84* deletion as heterozygous. We identified *let-7(n2853)-equivalent* mutants by singling F1 generation rollers and sequencing the non-Venus F2 generation progeny. The non-Venus quadruple mutants were invariably lethal at all temperatures that we tested but can be maintained by picking heterozygous animals expressing pharyngeal Venus.

We targeted *let-7* using a guide sequence overlapping with the mature *let-7* miRNA sequence and used ssODN repair templates to introduce single nucleotide blocking mutations into the endogenous *let-7* locus. As a proof of principle, to demonstrate our ability to create scarless edits within non protein-coding portions of the genome using a single nucleotide mismatch within the guide region, we aimed to generate a P6 blocking mutation, which recapitulates the classical *let-7(n2853)* hypomorphic mutation. *let-7(n2853)* disrupts the *let-7* seed sequence and leads to dysregulation of *let-7* mRNA targets in a temperature sensitive manner (Reinhart et al., 2000; Vella et al., 2004) (Figure 6B). We also designed two non-seed mutations as controls: one located within the PAM domain that is expected to completely block Cas9 and a second non-blocking mutation located downstream of the PAM (Figure 6B). We used *dpy-10* co-conversion to enrich for genome edited animals and singled both F1 generation roller and dumpy animals (Figure 6A). We then sequenced non-Venus F2 generation animals to examine how each blocking condition affected HDR efficiency (Figure 6A).

We observed high HDR-editing rates (60.7% HDR-edited) and low indel rates (10.7% indels) when the PAM was mutated (Figure 6C). We observed comparable rates of HDR-editing (56.7% HDR-edited) when P6, which recapitulates the *let-7(n2853)* mutation (Figure 6B), was used as the blocking mutation (Figure 6C). Although the HDR-editing rates were similar for the PAM and P6 blocking conditions, the indel rate was twice as high for the P6 blocking mutation (23.3% indels) compared to the PAM mutation (10.7% indels) (Figure 6C). The increased indel rate observed under P6 blocking conditions might indicate that P6 does not completely block Cas9. Consistent with this idea, we found that the no-blocking condition led to high frequency of indel mutations (40.7% indels) and low rate of HDR-editing (3.7% HDR-edited) (Figure 6C). Interestingly, we did not observe a similar increase in indel rates for the *tra-2* no-blocking condition (Figure 2, B-C), suggesting that there may be gene-specific or guide-specific differences in indel rates versus HDR-editing rates. Importantly, we were able to isolate a small percentage of *let-7* HDR-edited animals, even when the ssODN repair template did not contain a blocking mutation (Figure 6C). This observation supports the idea that in situations where no blocking mutations can be designed, desired edits can nonetheless be obtained, albeit at a low frequency.

Many microRNAs, including *let-7*, are members of microRNA families that share the same seed sequence and are therefore expected to target similar mRNA sequences (Lewis et al., 2003; Lim et al., 2003). As a result of sharing the same seed sequence, many members of microRNA families often exhibit functional redundancy with other family members (Miska et al., 2007; Alvarez-Saavedra and Horvitz, 2010; Abbott et al., 2005). As a further proof of principle, to demonstrate the utility of selecting for maternal chromosomes via paternally provided balancers, we aimed to recapitulate the *let-7(n2853)* variation in a genetic background devoid of the other three major *let-7* family members: miR-48, miR-84 and miR-241. Such strain has been difficult to generate using conventional genetic methods, since the *let-7* and miR-84 microRNAs are genetically linked and complete loss of *let-7* activity is lethal (Reinhart et al., 2000). To generate a strain containing mutations in all four *let-7* family members, we crossed *tmc24* balancer males to hermaphrodite animals containing deletions of *mir-48, mir-84 and mir-241* (Figure 6D). We then performed co-CRISPR of *let-7* and *dpy-10* to recapitulate the *let-7(n2853)* mutation (Figure 6D). As *let-7* and *mir-84* are both located on the X chromosome, F1 generation cross-progeny will have a paternally provided *tmc24* chromosome and a maternally provided chromosome that contains a mir-84 deletion, which we targeted for the *let-7* editing (Figure 6D). We then sequenced F2 generation non-Venus animals and identified animals containing maternal *let-7(n2853)* mutations. Using this strategy, we were able to create a stable strain that contains homozygous deletions in *mir-48* and *mir-241 (nDf51)* and the *mir-84(n4037)* deletion and *let-7(n2853*-equivalent) mutation balanced by *tmc24* in a heterozygous state.

Collectively, these findings demonstrate that single nucleotide substitutions within the guide RNA targeting sequence can be used to effectively mutate miRNAs through HDR in an otherwise scarless fashion. Furthermore, we propose that balancer chromosomes can be used to select for maternally provided haplotypes and introduce deleterious mutations directly into a balanced heterozygous background in a single injection step. The use of *dpy-10(cn64)* co-injection may further facilitate the single-step injection approach, since F1 generation rollers containing a single *dpy-10(cn64)* edited chromosome are more likely to only contain a single edit at a second locus. Previous strategies used in *C. elegans* to introduce potentially lethal genome edits directly into balanced genetic backgrounds have relied on two-step editing approaches, where the first edit introduces a nascent PAM to a gene of interest that can be specifically edited in a second injection after being crossed to a balancer chromosome that lacks the nascent PAM (Dejima et al., 2018; Duan et al., 2020). As nearly 90% of the *C. elegans* genome is covered by balancer mutations (Dejima et al., 2018), our strategy can be used to target most *C. elegans* genes in a single step editing approach, eliminating the need for extensive post-injection screening.

## Discussion

### Strategies for designing Cas9 blocking mutations in *C. elegans*

As effects of extraneous mutations are difficult to predict when editing non-protein coding genomic regions, we aimed to determine whether single nucleotide changes within the guide region of the donor were sufficient to promote HDR. Our findings confirm that mutating the PAM domain remains the most effective way to block Cas9 and leads to the highest HDR editing rates (Figure 7A and Figure 2). However, we demonstrate that single nucleotide substitutions in the guide sequence are also highly effective at blocking Cas9 and allow for excellent HDR editing rates. Therefore, when the desired mutation overlaps with the guide sequence, additional blocking mutations are not necessary to recover HDR edited animals, which can facilitate scarless HDR editing (Figure 7B and Figure 2). For protein coding genes, it is also possible to introduce silent blocking mutations into the guide sequence, ideally close to the 3’ end of the guide, when silently mutating the PAM is not possible (Figure 7C). Furthermore, we show that blocking mutations are not absolutely required for recovering HDR edited animals in *C. elegans*, although the HDR-editing efficiency is substantially reduced compared to when blocking mutations are used (Figure 7D and Figure 2). For non-coding regulatory sequence edits that do not overlap with a guide sequence, we suggest that it is possible to forego the use of blocking mutations in order to recover animals in an otherwise scarless genetic background (Figure 7D). The ability to recover HDR edited animals without including a blocking mutation suggests that a temporal block to Cas9 activity exists in *C. elegans,* preventing Cas9 recutting of the repaired genomic region (Figure 7E). Cas9 activity might be highest in the distal end of the maternal germline, since this is where Cas9 RNP complexes are injected (Figure 7E). The temporal block to Cas9 activity might result from degradation of the injected Cas9 RNP complexes or dilution of Cas9 activity as germ cells passage through the maternal germline from distal end to proximal end (Figure 7E). Collectively our findings expand the repertoire of possible genome edits in *C. elegans* and should facilitate analysis of non-coding regulatory sequences without the need for extraneous Cas9 blocking mutations.

**Figure 7.**
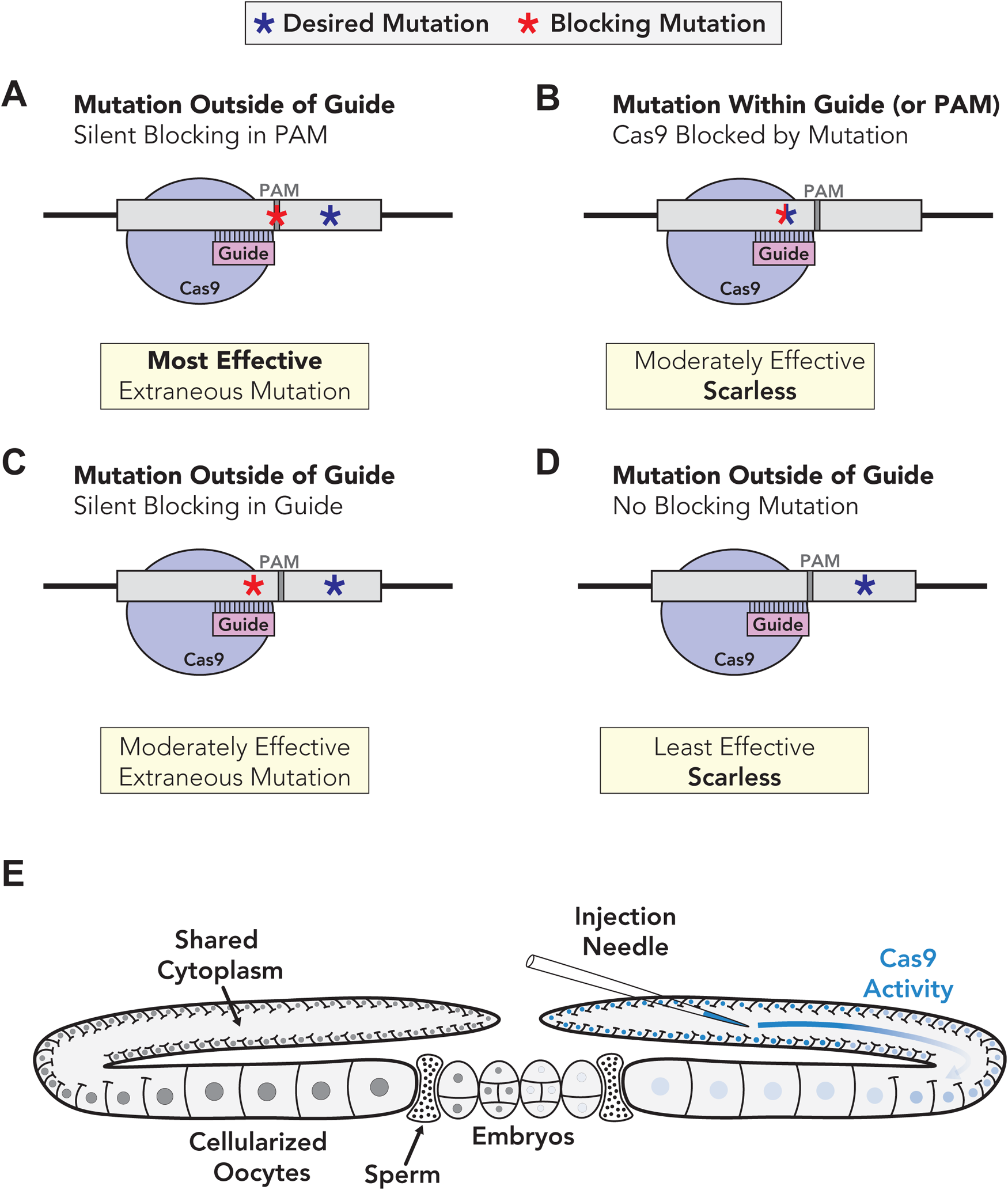
Strategies for creating Cas9 blocking mutations in *C. elegans.* (A-D) Examples of strategies for introducing mutations to block Cas9 in *C. elegans*. (A) The most straightforward method to blocking Cas9 is to introduce a silent mutation into the PAM domain. In an ideal scenario, the desired mutation disrupts the PAM domain and consequently blocks Cas9, which would lead to a “scarless” edit. Whenever possible, mutating the PAM completely blocks Cas9 and leads to the highest relative HDR rates. (B) When the intended mutation overlaps with the guide targeting sequence, additional blocking mutations are not required, although are slightly less effective than PAM mutations. An advantage of this method is that the desired mutation also serves as the blocking mutation, leading to a scarless edit. (C) For protein coding genes where the PAM cannot be silently mutated, a silent single base substitution in the guide targeting sequence is sufficient to promote HDR. (D) When silent mutations cannot be engineered into the guide or the desired mutation does not overlap with the guide/PAM sequences, it is possible to recover HDR-edited animals without including a blocking mutation in *C. elegans*. While this approach has the advantage of being scarless, it is the least effective in terms of HDR-editing rates. (E) Model for temporal block to genome editing in *C. elegans*. Cas9 RNPs and single-stranded DNA repair templates are injected into the distal end of the maternal germline (right gonad arm). Cas9 activity is attenuated as it passes from the injection site towards the proximal gonad. This might be due to two different possibilities: (1) Cas9 RNPs are unstable and/or degraded over time or (2) Cas9 is able to target chromosomes located in the distal germline more effectively than chromosomes in the proximal germline. In either case, Cas9 activity appears to be temporally restricted, allowing for HDR repair even when blocking conditions do not completely block Cas9.

### Differences in maternal vs. paternal genome editing rates in *C. elegans*

We found that the HDR editing rates of *tra-2* were much higher for the haplotypes that contained the *dpy-10(cn64)* allele compared to the haplotypes that were not edited for *dpy-10* (Figure 2). We hypothesized that the differences in HDR editing rates for each haplotype could reflect differences in the HDR editing rates for maternal and paternal haplotypes. Given that the Cas9 RNPs are injected into the maternal germline of hermaphrodite animals, it seemed likely that editing of the maternally provided haplotype is preferred over editing of the paternally provided haplotype. Indeed, several others have speculated that editing of maternal haplotypes is preferred in *C. elegans* (Paix et al., 2016; Kim et al., 2014; Arribere et al., 2014; Paix et al., 2014), although both paternal and maternal germ cells are competent for HDR (Clejan et al., 2006). To determine whether there was any preference for maternal or paternal editing, we used a mating-based approach that allowed us to quantify the editing rates of maternal and paternal haplotypes, demonstrating that maternal editing was preferred over paternal editing (Figure 5). This is in contrast to a recent study, which suggested that paternal genome (embryonic) editing is preferred over maternal editing (Farboud et al., 2018). A key difference in our experimental design was that we were able to assess maternal versus paternal genome editing in a single injection step, whereas previous studies have assessed maternal and paternal editing in separate injections (Farboud et al., 2018). Nevertheless, it is also possible that there could be gene-specific differences in haploid genome editing efficiency.

Taking advantage of the preference for maternal editing, we were able to select for the maternally provided chromosomes using balancer chromosomes that restrict recombination between the maternal and paternal chromosomes. By mating hermaphrodite animals to males containing a balancer chromosome before injection, the maternally provided non-balancer chromosome of co-edited F1 roller animals is more likely to be edited. Since typical balancer chromosomes contain a fluorescent or physical marker to identify animals harboring the balancer, non-marked animals can be easily identified and should be homozygous for the maternally provided haplotype.

Homozygosing for the edited, maternally provided chromosome would homozygose for the mutation of interest and therefore reduce molecular method-based screening. As most of the *C. elegans* genome is covered by balancer chromosomes (Dejima et al., 2018), this approach can be broadly applied to most *C. elegans* genes and has the added advantage of introducing potentially deleterious mutations directly into a balanced genetic background.

### Single nucleotide substitutions are sufficient to promote HDR in *C. elegans*

In this study, we demonstrate that a single nucleotide mismatch at any point in the guide sequence can block Cas9 and can be reliably used to recover HDR-edited animals.

Importantly, blocking Cas9 using single nucleotide substitutions helps minimize the number of extraneous mutations needed for genome editing experiments, which can facilitate scarless editing of genomic sequences that cannot be mutated in a silent manner. However, despite the fact that these single nucleotide blocking mutations are sufficient to promote HDR, we show that Cas9 was still able to target genomic sequences containing a single mismatch to its guide RNA, including the P2 mismatch located only 2 bases away from the 3’ end of the guide sequence (Figure 4). These data provide direct evidence that off-target cutting can occur in *C. elegans*, in contrast to previous speculation that off-target cutting may not readily occur in *C. elegans* (Schwartz and Sternberg, 2014; Xu, 2015). Previous targeted approaches to identify potential off-target cutting by Cas9 did not identify bona fide off-target events (Dickinson et al., 2013; Chiu et al., 2013; Silva-Garcia et al., 2019). Similarly, whole genome sequencing-based approaches did not identify variants at predicted off-target sites and the overall rate of variant formation was not significantly higher than the spontaneous mutation rate (Waaijers and Boxem, 2014; Au et al., 2018; Paix et al., 2014). Why haven’t previous approaches identified off-target cutting events in *C. elegans*? There are several contributing factors that might make identification of off-target cutting events difficult. We found that the efficiency of off-target cutting was much lower than on-target cutting (Figure 4), suggesting that off-target cutting events might be rare. Furthermore, off-target editing likely occurs in a heterozygous fashion, which can complicate detection in sequencing reactions since heterozygosity is rapidly lost in hermaphroditic organisms (Brenner, 1974). Another factor could be the software used to design guide RNAs, many (or all) of which might not allow guides that carry single mismatches to other regions of the genome. Although we did not test whether multiple guide substitutions were more effective at blocking Cas9, previous studies have suggested that increasing the number of mismatches in the guide sequence leads to increased blocking of Cas9 (Jinek et al., 2012; Fu et al., 2013; Pattanayak et al., 2013; Hsu et al., 2013; Mali et al., 2013; Cong et al., 2013; Zhang et al., 2015). Thus, effective guide RNA design might essentially eliminate off-target cutting by Cas9. Our observation that off-target cutting can occur in *C. elegans* emphasizes the importance of careful guide design and backcrossing of genome-edited strains to remove potential unwanted mutations when off-target effects are suspected.

### A temporal block to Cas9 activity appears to limit recutting of repaired DNA

We were surprised to find that we were able to recover HDR-edited animals when a blocking mutation was not introduced into ssODN repair templates, since Cas9 should still be able to target the repaired DNA. Although we observe significantly reduced HDR editing rates when no blocking mutation was included in the ssODN repair template, the editing efficiency was high enough that we were able to reliably recover HDR-edited animals. Given the ability of Cas9 to target double stranded repair templates, blocking conditions may remain critical for studies that use dsDNA repair templates. It is worth noting that many of the cases that might necessitate foregoing a blocking mutation, such as the editing of non-coding RNAs, would typically be small genomic changes that can be accomplished using ssODN repair templates.

The fact that we are able to recover HDR-edited animals without including a blocking mutation suggests that a temporal block to Cas9 activity exists in *C. elegans*, where the repaired region escapes repeated targeting by Cas9 (Figure 7E). What might lead to a temporal block to Cas9 activity? One possibility is that the reagents used for genome editing are not stable and/or targeted for degradation in the *C. elegans* germline, which would lead to reduced Cas9 activity over time. Consistent with this idea, RNP complexes are rapidly degraded (Prior et al., 2017; Kim et al., 2014; Liang et al., 2015; Farboud et al., 2018; DeWitt et al., 2017). Plasmid-expressed Cas9 may persist longer and might not result in the same temporal block that we observed for Cas9 RNP injection in this study. Interestingly, genome editing using double-stranded DNA templates and plasmid-expressed Cas9 was reported to still occur in progeny produced 64-90h post-injection (Ghanta and Mello et al., 2020). Thus, the choice of how Cas9 and repair templates are delivered may influence the length of time that Cas9 activity is permitted. A second explanation for a temporal block might be that germ cells could become less receptive to Cas9 as they passage through the *C. elegans* germ line. Cas9 RNP complexes are injected into the syncytial maternal germline, where immature germ cells share a common cytoplasm and can all be targeted by a single injection (Pazdernik and Schedl, 2012; Hubbard and Schedl, 2019; Evans et al., 2006; Kadandale et al., 2008). Many of the syncytial germ cells are in pachytene stage, during which time germ cells are receptive to homology directed repair pathways (Woglar and Jantsch, 2014, McClendon et al., 2016). As germ cells mature, they could become less amenable to genome editing. Lastly, it is also possible that genome editing reagents are diluted as they passage through the tubular-shaped maternal germline. This could explain the differences in editing efficiencies for maternal and paternal haplotypes, as the injection mix could become less available in the proximal germline. In any case, our findings suggest that Cas9 activity is attenuated over time, leading to a temporal block to Cas9 activity. This temporal block to Cas9 activity may then allow for effective HDR editing, even under conditions where Cas9 is not completely blocked. Additional work will be required to fully understand how a temporal block to Cas9 activity is established. It will also be interesting to see if a similar temporal block to Cas9 activity exists in other organisms, or if the unique germline architecture of *C. elegans* leads to reduced Cas9 activity over time.

## Acknowledgements

We thank current and former members of the Zinovyeva lab for valuable discussions and experimental assistance. We thank David Greenstein for providing helpful comments and suggestions. This work was supported by a grant from the National Institute of General Medical Sciences (R35GM124828 to A.Z.) and K-INBRE funds (P20 103GM418 to J.M. and J.S.) Some strains were provided by the CGC (*Caenorhabditis* Genetics Center), which is funded by the National Institutes of Health Office of Research Infrastructure Programs P40-OD010440.

## Supplementary Materials

**Table S1.** List of Strains Used in This Study.

| Strain | Genotype | Information |
| --- | --- | --- |
| FX30240 | *tmc24* [F23D12.4(*tmls1240*)] X | Dejima et al., 2018 |
| N2 | wildtype | From CGC |
| UY352 | *tra-2(zen142)* II | PAM + *Rsal* |
| UY356 | *tra-2(zen145)* II | *Rsal* only, no blocking |
| UY362 | *tra-2(zen151)* II | P2 + *Rsal* |
| UY364 | *tra-2(zen153)* II | P11 + *Rsal* |
| UY370 | *tra-2(zen157)* II | P20 + *Rsal* |
| UY386 | *let-7(zen162)/tmc24* X | PAM |
| UY389 | *let-7(zen165)/tmc24* X | No blocking |
| UY392 | *let-7(zen168)/tmc24* X | *let-7*(n2853-equivalent) |
| UY440 | *ndf51* V; *let-7(zen171)* *mir-84*(n4037)/*tmc24* X | *let-7* family mutant |
| VT1066 | *ndf51* V; *mir-84*(n4037) X | Abbott et al., 2005 |

**Table S2.**
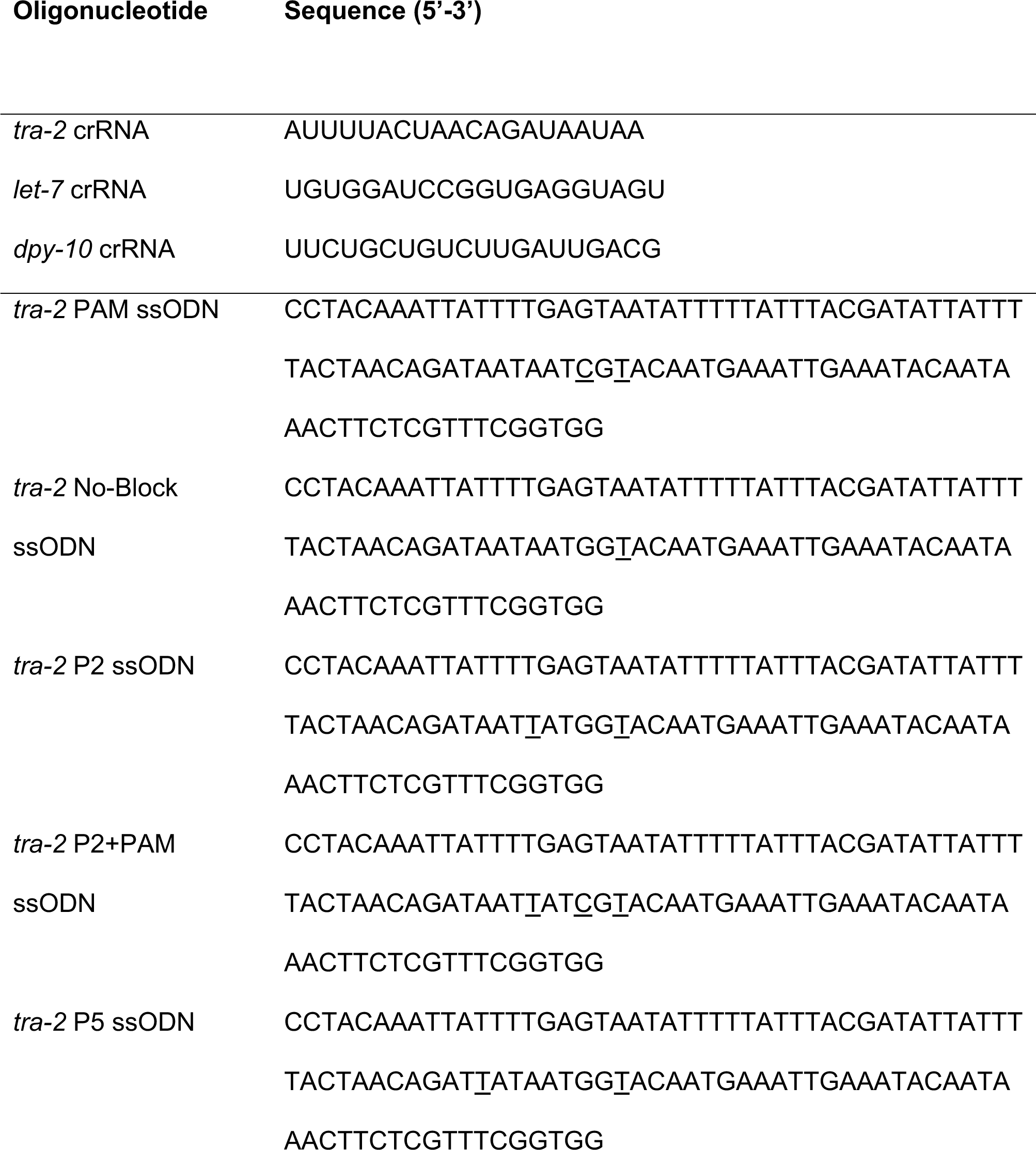
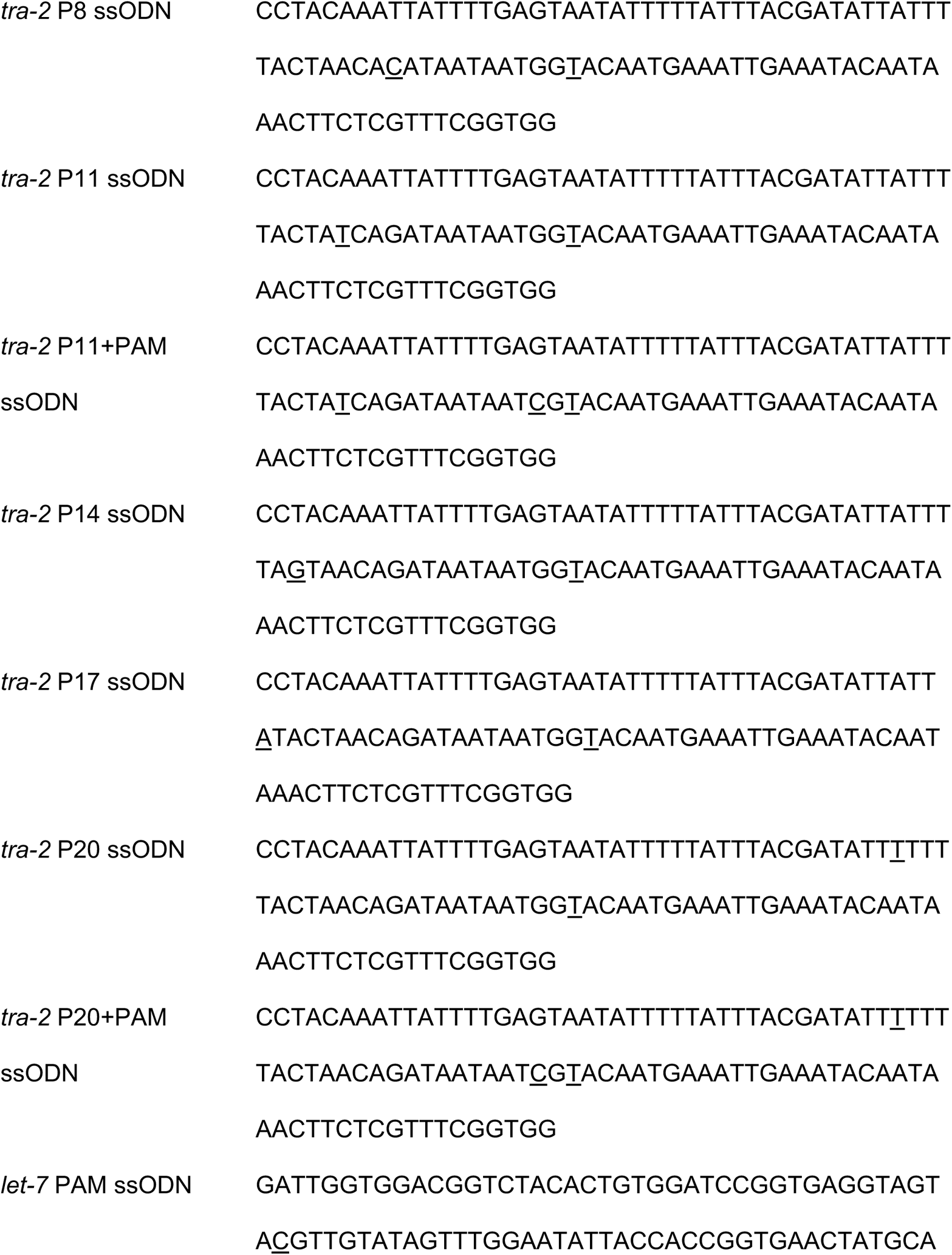
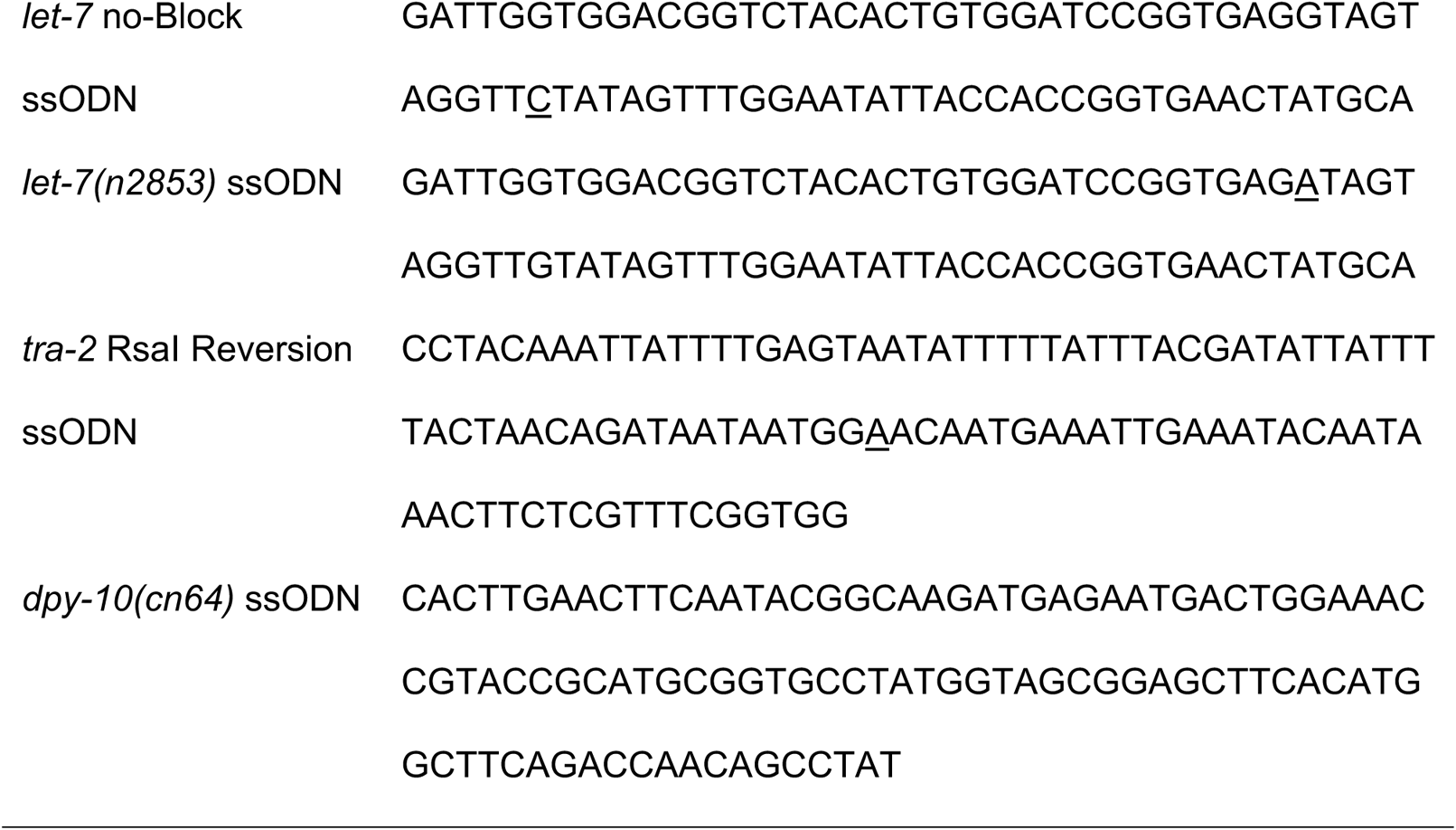
List of Oligonucleotides Used in This Study.

**Table S3.**
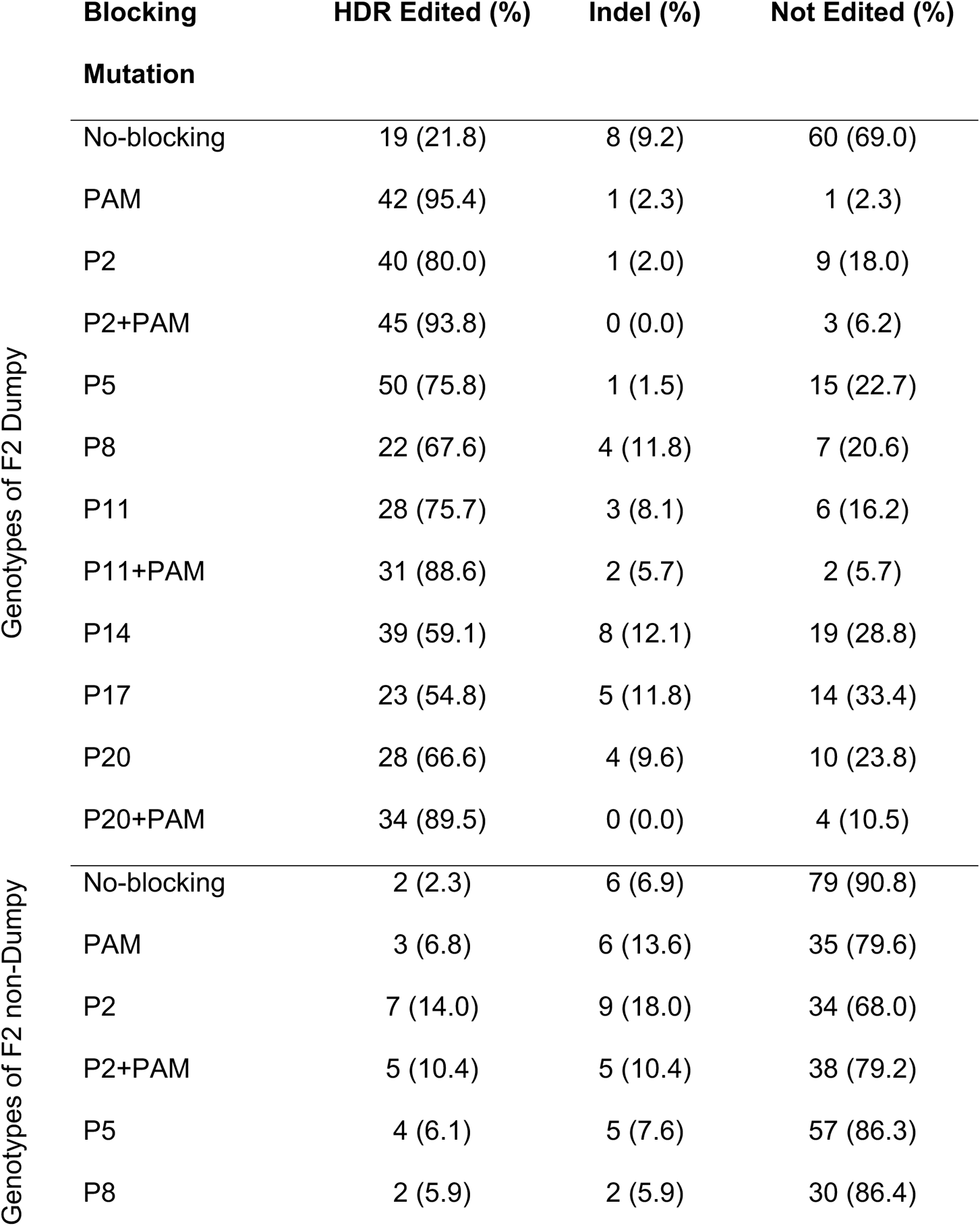
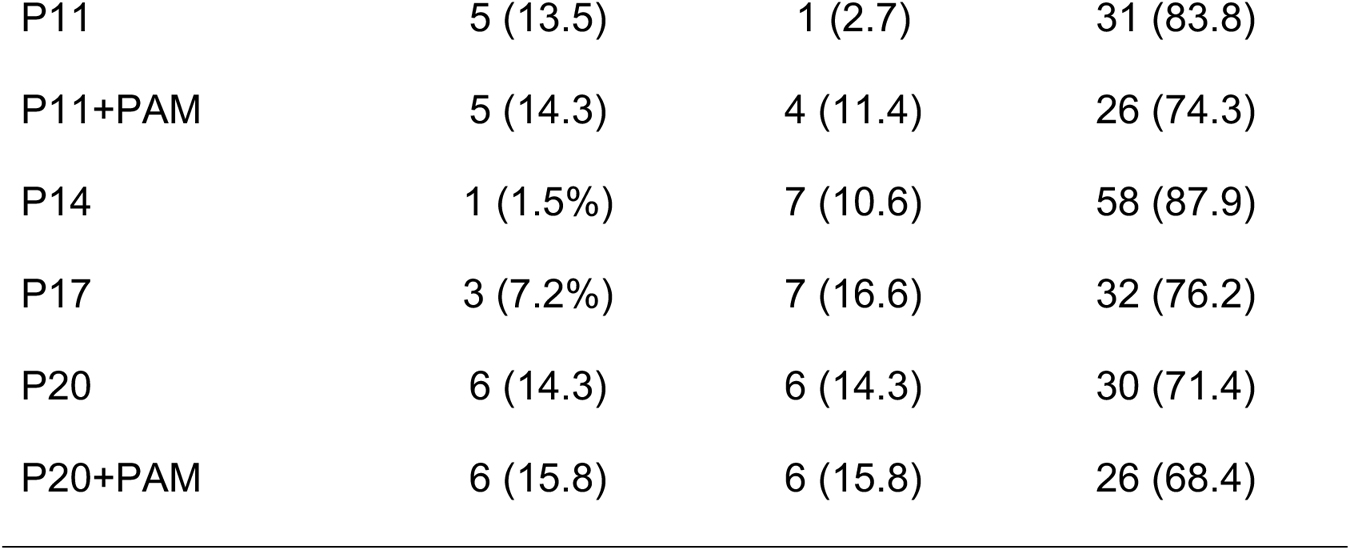
(Related to Figure 2). Editing Rates of *tra-2* in F2 Generation Dumpy (top) and Non-dumpy (bottom) Animals.

**Table S4.**
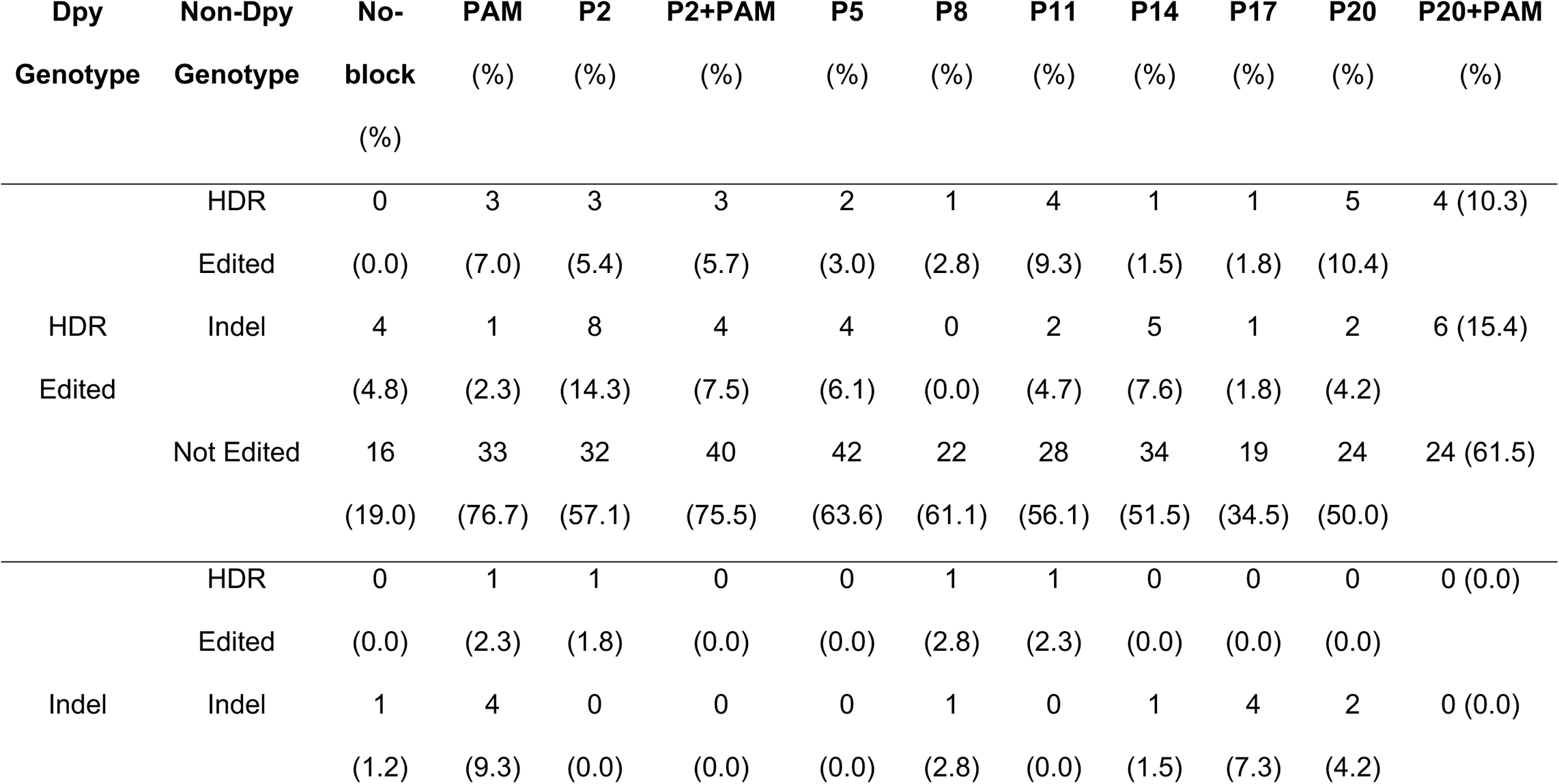
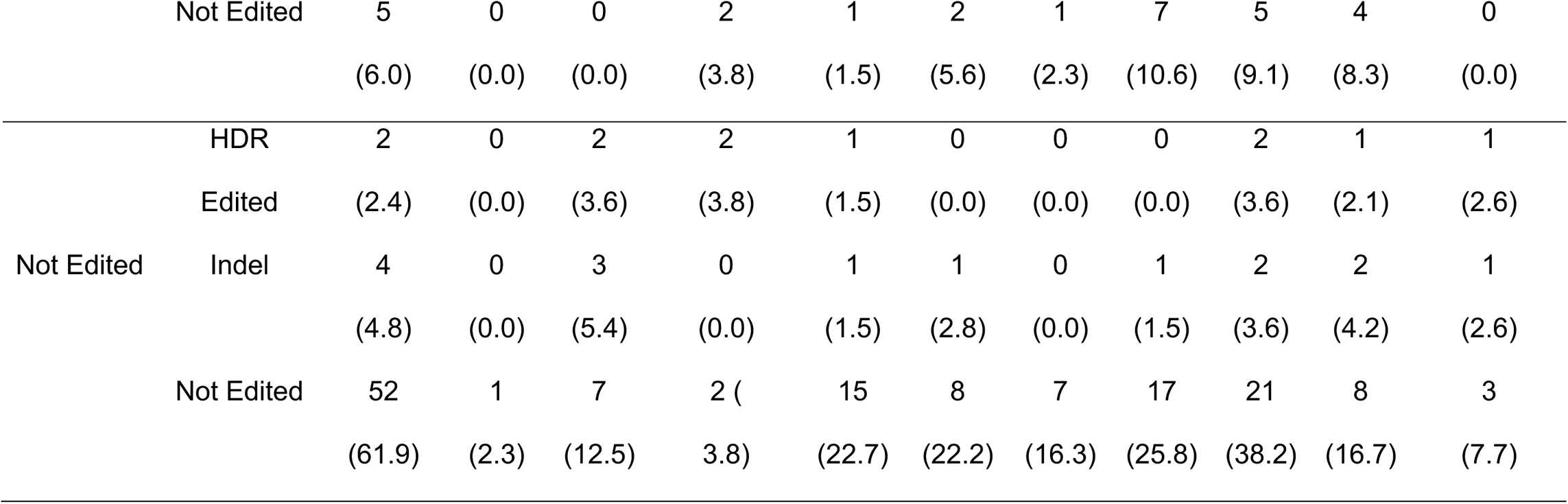
(Related to Figure 2D). Paired-Genotype Analysis of F2 Dumpy and Non-dumpy Animals from Single F1 Rollers.

**Table S5.** (Related to Figure 4A). HDR Incorporation Rates of Blocking Mutations and Non-blocking *Rsa*I Restriction Site Among HDR-edited Chromosomes.

| **Blocking Mutation** | ***RsaI* Only (%)** | **Blocking Only (%)** | **Blocking + *RsaI* (%)** |
| --- | --- | --- | --- |
| No-blocking | 21 (100.0) | 0 (0.0) | 0 (0.0) |
| PAM | 1 (2.2) | 0 (0.0) | 44 (97.8) |
| P2 | 15 (31.9) | 15 (31.9) | 17 (36.2) |
| P2+PAM | 0 (0.0) | 0 (0.0) | 52 (100.0) |
| P5 | 1 (1.8) | 40 (74.1) | 13 (24.1) |
| P8 | 2 (8.0) | 13 (56.0) | 8 (36.0) |
| P11 | 0 (0.0) | 21 (63.6) | 12 (36.4) |
| P11+PAM | 0 (0.0) | 10 (27.8) | 26 (72.2) |
| P14 | 8 (20.5) | 22 (56.4) | 9 (23.1) |
| P17 | 4 (15.4) | 16 (61.6) | 6 (23.0) |
| P20 | 4 (12.5) | 17 (53.1) | 11 (34.4) |
| P20+PAM | 0 (0.0) | 11 (28.2) | 28 (71.8) |

**Table S6.** (Related to Figure 3B). Effect of Distance to Cut Site on Incorporation of Single Nucleotide Guide Substitutions.

| **Blocking Mutation** | **PAM Only (%)** | **Blocking Only (%)** | **Blocking + *RsaI* (%)** |
| --- | --- | --- | --- |
| PAM | 45 (100.0) | 0 (0.0) | 0 (0.0) |
| P2+PAM | 0 (0.0) | 0 (0.0) | 52 (100.0) |
| P11+PAM | 26 (72.2) | 4 (11.2) | 6 (16.6) |
| P20+PAM | 28 (71.8) | 0 (0.0) | 11 (28.2) |

**Table S7.** (Related to Figure 4). Blocking Efficacy of Single Nucleotide Substitutions.

| Blocking Mutation |  | Both Reverted (%) | Heterozygous (%) | Not Reverted (%) |
| --- | --- | --- | --- | --- |
| F1 RoI | No-blocking | 0 (0.0) | 25 (50.0) | 25 (50.0) |
|  | PAM | 0 (0.0) | 0 (0.0) | 56 (100.0) |
|  | P2 | 0 (0.0) | 1 (1.8) | 56 (97.2) |
|  | P11 | 0 (0.0) | 3 (5.4) | 53 (94.6) |
|  | P20 | 0 (0.0) | 7 (12.7) | 48 (87.3) |
| F1 Dpy | No-blocking | 5 (10.0) | 22 (44.0) | 23 (46.0) |
|  | PAM | 0 (0.0) | 0 (0.0) | 50 (100.0) |
|  | P2 | 0 (0.0) | 3 (5.3) | 53 (94.7) |
|  | P11 | 0 (0.0) | 3 (5.3) | 53 (94.7) |
|  | P20 | 1 (1.8) | 12 (21.8) | 42 (76.4) |

**Table S8.** (Related to Figure 6). Editing Rates of let-7 in F2 Generation non-Venus Animals.

| **Blocking Mutation** | **HDR Edited (%)** | **Indel (%)** | **Not Edited (%)** |
| --- | --- | --- | --- |
| No-blocking | 1 (3.7) | 11 (40.7) | 15 (55.6) |
| PAM | 17 (60.7) | 3 (10.7) | 8 (28.6) |
| *n*2853 [P6] | 17 (56.7) | 7 (23.3) | 6 (20.0) |

